# Repurposing the TIGR-Tas system for programmable transcription activation

**DOI:** 10.64898/2026.07.30.739400

**Authors:** Yetong Sang, Lingjie Xu, Nga Man Wong, Yongfei Chang, Zhenkun Cai, Zehua Bao

## Abstract

Programmable transcription activators are central to functional genomics, gene therapy, and synthetic biology. However, application of the most versatile CRISPR activators (CRISPRa) is hampered by the bulky size and compact CRISPR systems are constrained by the requirement of long protospacer adjacent motifs (PAMs). To address these limitations, we repurposed the TIGR-Tas RNA-guided DNA-targeting system as a versatile platform for gene activation. By systematically optimizing the fusion architecture of a nuclease-inactivated TasR with gene activation domains, we achieved robust, specific, and multiplexable transcriptional activation of endogenous human genes. This TIGR activation (TIGRa) system could be a superior alternative to CRISPRa systems due to its extreme compactness and potentially PAM-independent targeting capability.

## INTRODUCTION

Artificial transcription factors enable modulation of gene expression without altering genomic sequence and have transformed functional genomics, synthetic biology, and therapeutics^1–5^. Notably, CRISPR-based activation (CRISPRa) systems fuse nuclease-dead Cas proteins to transcriptional effectors, providing programmable transcriptional control of endogenous loci by specifying the guide RNA (gRNA) sequence^6,7^. Diverse activator architectures, including multi-domain fusions such as SAM, SunTag, and VPR, achieved high efficiency gene activation^8–11^. In parallel, alternative CRISPR systems were repurposed to expand genomic targeting range^12^.

Despite these advances, nearly all Cas proteins require protospacer adjacent motifs (PAMs) for DNA targeting, which constrain targetable sequences and limit dense regulatory tiling^13^. In addition, commonly used Cas proteins are large in size and complicate delivery. The recent discovery of TIGR-Tas systems offers a compelling alternative. TIGR-Tas systems comprise an extremely compact effector protein, Tas, that binds short gRNAs encoded in TIGR arrays^14^. In contrast to CRISPR, TIGR-Tas uses a tandem interspaced gRNA (tigRNA) mechanism without an obligatory PAM, potentially enabling PAM-free targeting^14^. Here, we repurposed the TasR nuclease for programmable transcriptional activation by converting it into a catalytically dead scaffold and enhanced its potency through the SunTag strategy. We show that TIGR activation (TIGRa) is highly specific and potentially multiplexable.

## RESULTS

### Turning TIGR-Tas into programmable transcription activators

We started with two Tas proteins, TasA and *Par*TasR. TasA naturally lacks a nuclease domain and thus only binds DNA without cleaving, serving as a natural DNA-binding platform. *Par*TasR contains an N-terminal RuvC nuclease domain and was shown to induce indels in mammalian cells. To convert *Par*TasR into a DNA-binding platform, we made nuclease-inactivated versions of *Par*TasR by either mutating the metal-binding aspartate residue (D7) to an alanine residue (**Supplementary Fig. 1**, referred to as dTasR) or truncating the full RuvC domain (**Supplementary Fig. 2**, referred to as tTasR). We then designed and evaluated a panel of TasA- and TasR-based activators (**Fig. 1, Supplementary Fig. 3**, and **Supplementary Table 1**).

**Figure 1.**
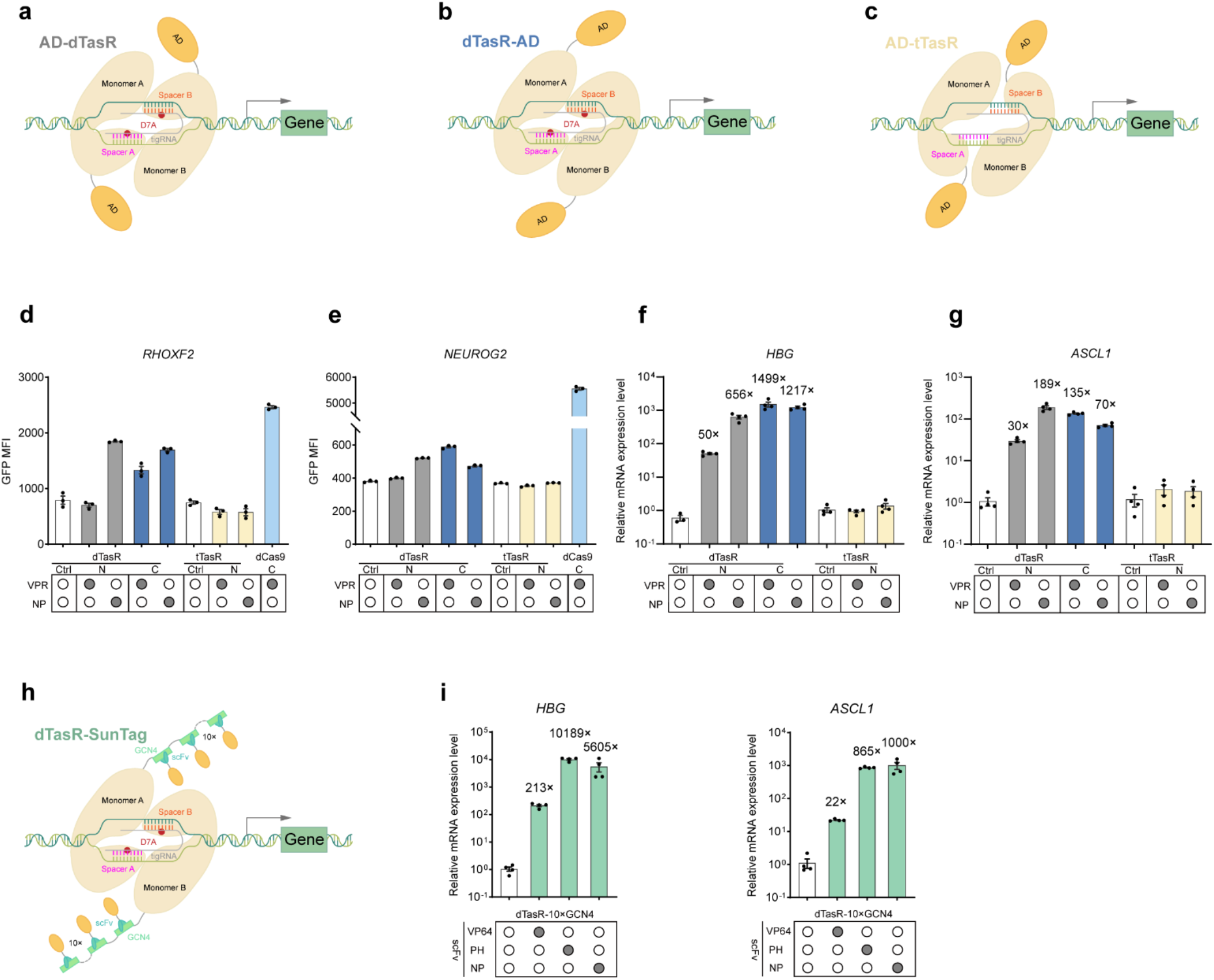
TIGRa-mediated transcription activation of endogenous human genes. **a**-**c**, Schematic of N-terminally fused AD-dTasR (**a**), C-terminally fused dTasR-AD (**b**), and N-terminally fused AD-tTasR (**c**). **d**-**e**, The EGFP mean fluorescence intensity induced with different dTasR and tTasR fusions at *RHOXF2-EGFP* reporter locus (**d**) and *NEUROG2-EGFP* reporter locus (**e**). Ctrl, dTasR or tTasR without AD. The data were graphed as mean ± S.E.M and represent three biological replicates. **f**-**g**, Relative expression levels of different dTasR and tTasR fusions at *HBG* (**f**) and *ASCL1*(**g**) in HEK293T. The housekeeping gene *glyceraldehyde phosphate dehydrogenase* (GAPDH) was used as an internal control for the normalization of qRT-PCR data. Ctrl, dTasR or tTasR without AD. The data were graphed as mean ± S.E.M and represent four biological replicates. **h**, Schematic of SunTag platform based on dTasR. **i**, Relative expression levels of dTasR-SunTag systems at *HBG* and *ASCL1* in HEK293T. The housekeeping gene *glyceraldehyde phosphate dehydrogenase* (GAPDH) was used as an internal control for the normalization of qRT-PCR data. Ctrl, dTasR-SunTag without AD. The data were graphed as mean ± S.E.M and represent four biological replicates.

Transcriptional activation activity was benchmarked at protein level in HEK293T cells using green fluorescent protein reporters at *RHOXF2* and *NEUROG2* loci (*RHOXF2*-EGFP and *NEUROG2*-EGFP fusions, **Supplementary Fig. 4** and **Supplementary Table 2**) for primary evaluation, followed by validation at mRNA level at two additional endogenous loci (*HBG* and *ASCL1*) to confirm functionality on natural genomic targets (**Fig. 1**). For each target gene, we designed four tigRNAs by selecting an 18 bp target sequence within previously validated functional Cas9 gRNA protospacer sequences (**Supplementary Table 3**)^6,10^. Reporter cell lines were co-transfected with plasmids expressing Tas-AD variants and tigRNA pools.

We generated direct protein fusions with potent activation domains (ADs), including VPR and NP to the N or C terminus of Tas proteins^15,16^ (**Fig. 1a,b** and **Supplementary Fig. 3a**). For tTasR, we fused ADs to its N-terminus in replacement of the RuvC domain, to investigate the contributions of RuvC for DNA-binding (**Fig. 1c** and **Supplementary Fig. 3a**). For direct TasA-AD fusions, we did not observe gene activation at two EGFP reporter loci (**Supplementary Fig. 5a**). At *HBG* and *ASCL1*, TasA-VPR induced observable but low activation. This activation was amplified at *HBG* by using TasA-NP (**Supplementary Fig. 5b**). TasA-SunTag fusions also induced activation, especially at *ASCL1* (**Supplementary Fig. 5c**).

However, activation levels induced by TasA were generally low (less than 100-fold). In contrast, direct dTasR-VPR (879 aa, **Supplementary Table S1**) and dTasR-NP (808 aa, **Supplementary Table S1**) fusions yielded robust activation at EGFP reporter loci and the two endogenous genes (**Fig. 1d-g, Supplementary Fig. 6**, and **Supplementary Fig. 7**), inducing 70-to-1499-fold activation over base expression level at *ASCL1* and *HBG* (**Fig. 1f** and **g**). The more compact dTasR-NP produced similar or slightly lower activation depending on the locus (**Fig. 1d-g**). The N-terminal fusion NP-dTasR showed the highest activation at *RHOXF2*-EGFP reporter locus and *ASCL1* (**Fig. 1d** and **g**), whereas VPR-dTasR showed substantially reduced activation, suggesting that VPR is more compatible as a C-terminal fusion. Using an extended XTEN80 linker between dTasR and AD reduced activation in all cases (**Supplementary Fig. 8**). In addition, replacing the RuvC domain with an AD completely abolished gene activation, suggesting that the RuvC domain contributes to DNA binding (**Fig. 1d-g**).

### Improving activation potency through signal amplification

To further improve activation potency, we sought to increase the copy number of ADs recruited to dTas. We first implemented MS2-mediated AD recruitment by fusing an MS2 hairpin to either the 5’ or 3’ end of the tigRNA, which we hypothesize will minimally affect the tigRNA structure as compared to internal insertions, and co-expressing MCP-p65HSF1 fusions^8,17^ (**Supplementary Fig. 3b** and **Supplementary Fig. 9a**). For dTasR, we observed low activation when the MS2 loop was fused to the 3’ end of the tigRNAs, which was observed at *ASCL1* and *HBG* sites (**Supplementary Fig. 9b**,**c**). MS2-based recruitment using TasA again induced no observable activation, consistent with TasA direct fusions (**Supplementary Fig. 10**). In light of this low activation potency as compared to direct protein fusions, we suspect that tigRNAs may be sensitive to structural perturbations.

We then employed the SunTag signal amplification system by fusing an array of GCN4 peptide repeats to the C-terminus of Tas proteins for recruiting multiple copies of scFv-AD fusions^9^ (**Fig.1h** and **Supplementary Fig. 3c**).We tested dTasR-based SunTag systems. dTasR-SunTag with 10×GCN4 repeats combined with scFv–PH or scFv–NP achieved high activation efficiencies, ranging from 865 to 10188-fold (**Fig. 1i**). This represents an order-of-magnitude improvement than direct AD fusions (**Fig. 1f** and **g**). The above results indicate that dTasR C-terminal fusions and dTasR-SunTag represent potent designs of TIGR-Tas based gene activation (referred to as TIGRa hereafter).

### Pooled tigRNAs show synergistic effects

Previous CRISPRa studies have revealed synergistic effects among CRISPR gRNAs^7^, where pooled gRNAs induced higher transcriptional activation than the additive effect of individual gRNAs. To evaluate whether tigRNAs can exhibit similar synergistic effects, we compared the activation potency of individual tigRNAs against pooled tigRNAs using dTasR-VPR. Indeed, pooled tigRNAs yielded synergistic activation at *RHOXF2, NEUROG2*, and *HBG*, exceeding the sum of individual tigRNAs (**Fig. 2a,b** and **Supplementary Fig. 11**).

**Figure 2.**
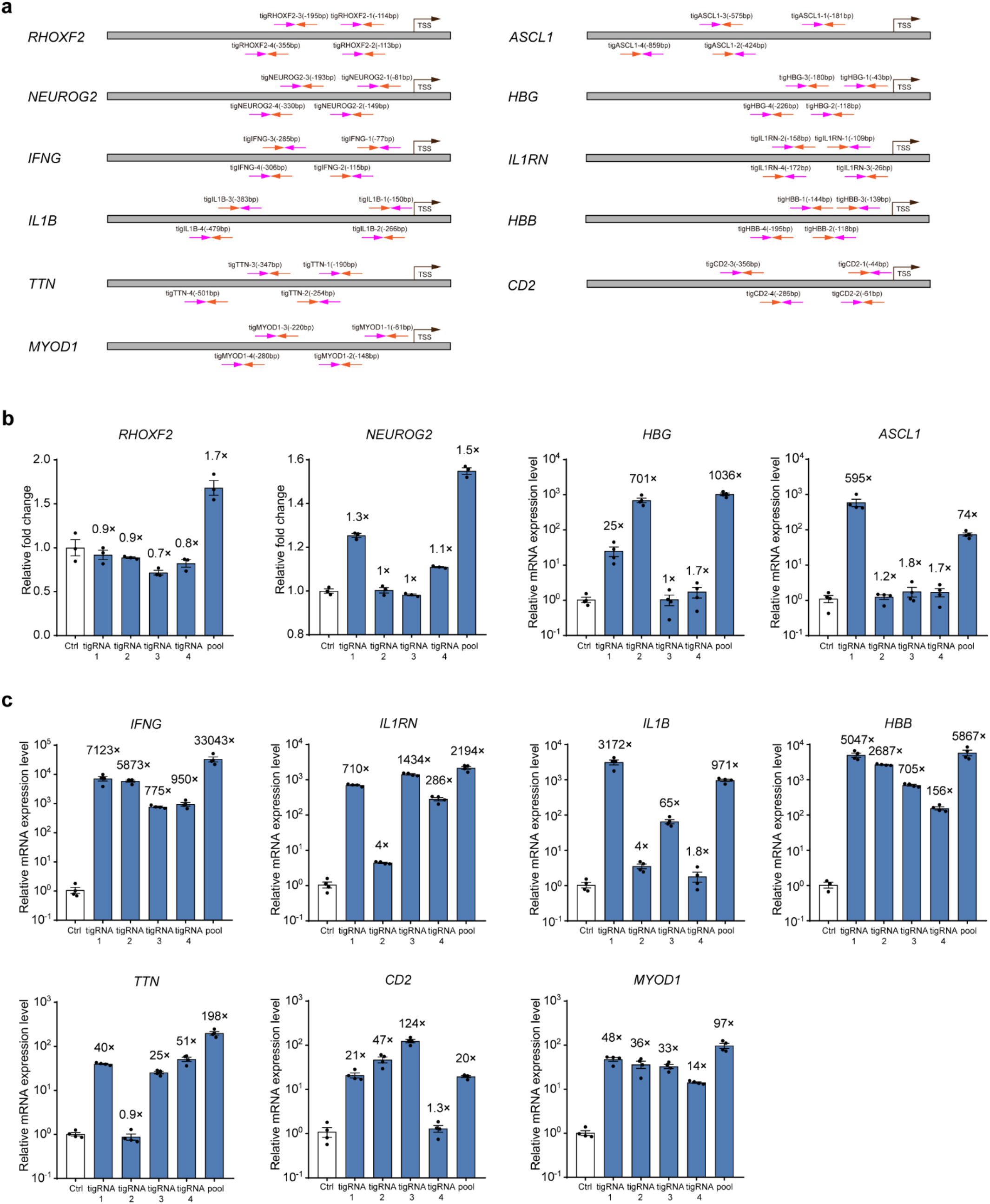
tigRNA synergy of TIGRa. **a**, Location of tigRNA target sequences of TasR for endogenous gene activation. The magenta arrows indicate spacer A and the orange arrows indicate spacer B. **b**, Activation of two EGFP reporter loci and endogenous human genes with dTasR-VPR using either individual or pooled tigRNAs. Ctrl in *RHOXF2*-GFP and *NEUROG2*-GFP loci is dTasR without AD. The data were graphed as mean ± S.E.M and represent three biological replicates. Ctrl in *HBG* and *ASCL1* experiments is dTasR-VPR without tigRNAs. The data were graphed as mean ± S.E.M and represent four biological replicates. **c**, Activation of endogenous human genes with dTasR-SunTag-PH using either individual or pooled tigRNAs. Ctrl, dTasR-SunTag-PH without tigRNAs. The data were graphed as mean ± S.E.M and represent at least three biological replicates.

To further demonstrate the versatility of TIGRa at more loci, we tested the activation of dTasR-SunTag-PH at seven additional endogenous genes (*IFNG, IL1RN, IL1B, HBB, TTN, CD2* and *MYOD1*, **Fig. 2c**). Adopting the same design principles, we designed four tigRNAs for each gene within previously used Cas protein gRNA protospacer sequences (**Supplementary Table 3**)^8,10,18–22^. All seven genes were activated using pooled tigRNAs. Among 28 individual tigRNAs, 23 showed activation with over 10-fold transcription improvement. Synergistic activation by pooled tigRNAs was observed at *IFNG* and *TTN* loci, especially for *IFNG* (up to 33043-fold, **Fig. 2c**). The observed synergy is consistent with CRISPRa experiments and suggests that tigRNA pools can be optimized to further improve the overall activation efficiency.

### TIGR array–mediated multiplex gene activation

Leveraging the selected tigRNAs, we engineered two customized TIGR arrays harboring triple tandem-linked tigRNAs targeting *NEUROG2, ASCL1* and *HBG*, or alternatively *ASCL1, HBG*, and *IFNG* using the most efficient tigRNA for each gene (**Fig. 2b,c** and **Fig. 3a,b**). Using dTasR-SunTag systems, both TIGR arrays effectively induced transcriptional activation of their respective target genes. For *NEUROG2, ASCL1* and *HBG* target genes, TIGR array–based activation elicited markedly greater transcriptional enhancement compared with pooled tigRNAs, yielding activation levels ranging from 2 to 7802-fold (**Fig. 3a** and **Supplementary Fig. 12**). When targeting *ASCL1, HBG*, and *IFNG* genes, dTasR-SunTag-NP induced transcriptional activation ranging from 86 to 581-fold, whereas dTasR-SunTag-PH elicited stronger activation, reaching 131 to 1314-fold induction.

**Figure 3.**
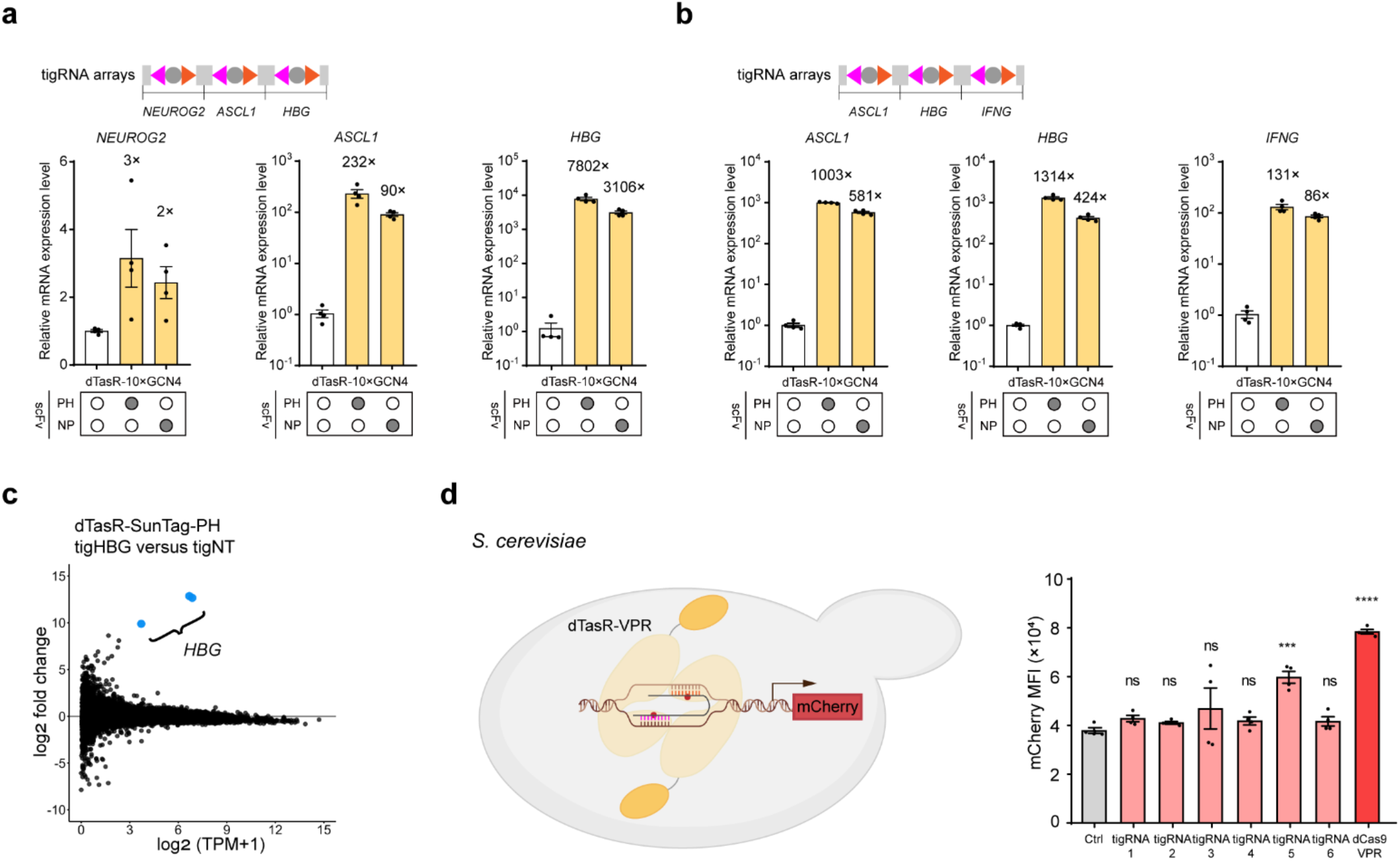
Multiplex gene activation, gene activation specificity and TIGRa performance in yeast. **a**, Multiplex activation of three endogenous genes *NEUROG2, ASCL1* and *HBG* with dTasR-SunTag using TIGR array. Ctrl, dTasR-SunTag without AD. The housekeeping gene *glyceraldehyde phosphate dehydrogenase* (GAPDH) was used as an internal control for the normalization of qRT-PCR data. The data were graphed as mean ± S.E.M and represent four biological replicates. **b**, Multiplex activation of three endogenous genes *ASCL1, HBG* and *IFNG* with dTasR-SunTag using TIGR array. Ctrl, dTasR-SunTag without AD. The data were graphed as mean ± S.E.M and represent four biological replicates. **c**, Transcriptome profiling of dTasR-SunTag-PH targeted to the *HBG* locus. HEK293T cells transiently expressing dTasR-SunTag-PH together with *HBG*-targeting tigRNAs were harvested 72 h post-transfection for RNA-seq analysis. The data represent the mean of three biological replicates. TPM, transcripts per kilobase million. **d**, Schematic of dTasR-VPR activation in yeast and mCherry activation levels of dTasR-VPR with different tigRNAs targeting *CYC1* promoter in yeast. Ctrl, dTasR-VPR without tigRNA. The data were graphed as mean ± S.E.M and represent four biological replicates. Significance levels were calculated by one-way ANOVA followed by Dunnett’s test against Ctrl. ***, P < 0.001; ****, P < 0.0001; ns, not significant.

### TIGR activation is highly specific

Having identified dTasR-SunTag-PH as the best-performing fusion architecture for potent activation, we assessed its genome-wide specificity profile. We performed RNA-seq on HEK293T cells expressing *HBG*-targeting dTasR-SunTag-PH. We found that the correlation in gene expression between *HBG*-targeting dTasR-SunTag-PH and control samples with non-targeting tigRNAs was very similar to the correlation between biological replicates in our data set (R^2^ > 0.97 in each case), indicating that gene expression is not broadly affected by the genomic targeting of dTasR-SunTag-PH (**Supplementary Fig. 13**). Disregarding noise from genes with low basal expression, *HBG* was the most highly upregulated gene, indicating that dTasR-SunTag-PH is highly specific (**Fig. 2c**).

### TIGR activation is portable to yeast

Finally, to assess its applicability beyond human cells, we examined TIGRa portability in *Saccharomyces cerevisiae* (**Fig. 2d**). dTasR-VPR was constitutively expressed and tigRNAs targeting the CYC1p promoter of a genomically integrated mCherry reporter were expressed from a small RNA promoter SNR52p^23,24^. The results showed that dTasR-VPR significantly improved mCherry expression in yeast when guided by tigRNA 5 (**Fig. 2d**). Thus, TIGRa is portable to the single cell model eukaryote and may be functionally extended to other species.

## DISCUSSION

We have leveraged the modularity of the TIGR-Tas RNA-guided DNA-targeting system to develop programmable transcriptional activators in eukaryotic cells. dTasR direct fusions and SunTag systems achieved robust transcriptional activation. Furthermore, TIGR activation is highly specific and multiplexable. The compact nature of TIGRa makes this system well-suited for viral delivery and translational applications. Future work is needed to define tigRNA design rules and potentially develop genome-scale, PAM-free tigRNA libraries for high-resolution functional genomics. With currently observed efficiency as a starting point, further Tas protein engineering offers potential paths to improve gene activation potency, potentially surpassing CRISPRa systems.

## ACKNOWLEDGEMENTS

This work was supported by the National Key R&D Program of China (grant no. 2023YFF1204500 to ZB), the National Natural Science Foundation of China (grant no. 22308316 to ZB), and the Fundamental Research Funds for the Central Universities (grant no. 226-2025-00043 to ZB). We thank iBioFoundry and Core Facility at ZJU‐Hangzhou Global Scientific and Technological Innovation Center for their technical support.

## AUTHOR CONTRIBUTIONS

Z.B. conceived of the study. Y.S., L.X., Y.C., N.M.W., and Z.C. constructed all the plasmids. Y.S. performed the endogenous gene activation experiments. L.X. performed the GFP-tagged *RHOXF2* and *NEUROG2* activation experiments. Y.S. and L.X. performed the RNAseq experiment. Y.S. and Z.C. performed activation experiments in yeast. Y.S. and L.X. analyzed the data. Y.S., L.X., and N.M.W. wrote the manuscript and made the figures. Z.B. reviewed and edited the manuscript. All authors read and approved the final manuscript.

## COMPETING FINANCIAL INTERESTS

Z.B., Y.S., L.X., Y.C., and N.M.W. are inventors on a patent application submitted by Zhejiang University that covers the TIGR activation technology described in this study.

## ADDITIONAL INFORMATION

Supplementary information is available in the online version of the paper. Correspondence and requests for materials should be addressed to Z.B.

## METHODS

### Plasmid construction

Plasmids were cloned by standard molecular cloning techniques. FpTasA, ParTasR, MCP, and XTEN80 gene fragments were human codon-optimized and commercially synthesized (Jiutian Gene Technology, Tianjin, China). GCN4 and scFv gene fragments were human codon-optimized and commercially synthesized (GenScript, China). pZB-TasA-p2A-BFP and pZB-dTasR-p2A-BFP (derived form pZB-dCas9-p2A-BFP backbone by replacing dCas9 with TasA or dTasR, respectively) were used as the backbone for TasA- and dTasR-TAD fusion constructs. dTasR was generated by mutating D7 to A7 through PCR-mediated site-directed mutagenesis. tTasR was generated by deleting RuvC domain (M1 to G100) of ParTasR. VP64 and VPR gene fragments were amplified from pRS415-Cas9-VPR (Addgene #163971). NP gene fragment was amplified from pZB-dCas9-NP. P65-HSF1 gene fragment was amplified from pAC1410 and ligated to pZB-MCP-p2A-BFP backbone. tigRNA expression plasmids were constructed by ligating the corresponding annealed oligos to the backbone plasmid pGL3-U6-tigRNA-BSD (derived by replacing the EGFP gene of Addgene #107721 with a blasticidin S deaminase gene marker and replacing Cas9 sgRNA scaffold with TasA or TasR tigRNA scaffold) downstream of the human U6 promoter. The plasmids used for yeast mCherry activation were constructed using pCRCT (Addgene #60621) as the backbone. The Cas9 fragment was replaced by dTasR. The *URA3* selection marker was replaced by *LEU2*, which was amplified from p415-GalL-Cas9-CYC1t (Addgene #43804). The VPR fragments were amplified from corresponding pZB-dCas9-VPR plasmids and inserted between the NLS at the C-terminus of dTasR and the ADH2 terminator. The tigRNA sequences were cloned into the backbone of each pCRCT-dTasR-VPR plasmid by amplifying the whole plasmid with tigRNA sequences as part of two primers. The sequences of all tigRNAs are listed in **Supplementary Table S3**.

### Cell culture

HEK293T (Cat. # GNHu44) cells were purchased from the National Collection of Authenticated Cell Cultures (Shanghai, China) and cultured in DMEM with high glucose additionally supplemented with 10% FBS (ExCell Bio, China), GlutaMAX (Gibco, USA) and MEM non-essential amino acids (Gibco, USA). Cells were grown at 37 °C under 5% CO_2_ in a humidified incubator and maintained at confluency below 90%.

### Generation of the EGFP-tagged endogenous reporter cell lines

Endogenous genes *RHOXF2* and *NEUROG2* were selected as the target sites for reporter cell lines constructions. pMP472-D2 encoding an enhanced GFP (EGFP) was used as homology directed repair donor plasmid. pMP472-D2 donor plasmid was derived from pMP472 (Addgene #134997) by deleting pMinCMV and pGK-BGH pA fragment, and replacing ZF binding sites with corresponding endogenous gene homology arms by Gibson assembly. Homology arms of gene *RHOXF2* and *NEUROG2* were amplified from HEK293T genomic DNA. The endogenous gene targeting CRISPR/Cas9 plasmids were constructed by inserting sgRNA sequence into pLentiCRISPRv2-neo backbone (Addgene #127644, a gift from Dr. Xia Liu, Zhejiang University) by Golden Gate assembly. Sequences of sgRNAs for CRISPR/Cas9 mediated-HDR are listed in **Supplementary Table S2**. The donor plasmid was co-transfected with corresponding pLentiCRISPRv2-neo targeting plasmid into HEK293T to carry out EGFP chromosomal integration. 3 days post-transfection, cells were dissociated and 8×10^5^ cells were plated into 6-well plates. 1 day after plating, cells were transfected again with pZB-dCas9-VPR and corresponding pooled sgRNAs to activate EGFP expression. 3 days after the secondary transfection, GFP-positive monoclonal cells were sorted using BD FACSAria™ III (USA). Monoclonal cells were initially validated using flow cytometry. Only those showing activated GFP peak after transcriptional activation were chosen.

The correct insertion of EGFP sequence into endogenous gene loci was validated by genomic PCR.

### Transfection

All HEK293T cells were transfected with polyethyleneimine (PEI, YEASEN, China). For qRT-PCR assay in wild type HEK293T, the total amount of plasmids was 500 ng per well for a 24-well plate. Approximately 2×10^5^ cells were plated per well one day before transfection. For TasA- and dTasR-TAD direct fusions experiments, 250 ng plasmids encoding TasA-TAD or dTasR-TAD, and 250 ng 4 pooled tigRNAs were co-transfected. For TasA- and dTasR-SAM, -Suntag experiments, 166 ng TasA or dTasR, 166 ng secondary TAD fusion components (i.e. MCP-P65-HSF1, scFv-VP64, scFv-P65-HSF1, scFv-NP, respectively) and 166 ng 4 pooled tigRNAs plasmids were used. For flow cytometry analysis in EGFP reporter cell lines, the total amount of plasmids was 600 ng per well for a 24-well plate. For TasA- and dTasR-TAD direct fusions experiments, 400 ng plasmids encoding TasA-TAD or dTasR-TAD, and 200 ng 4 pooled tigRNAs were co-transfected. For TasA- and dTasR-SAM experiments, 200 ng TasA or dTasR, 200 ng MCP-p65-HSF1 and 200 ng 4 pooled tigRNAs plasmids were used. 24 hours after transfection, the culture medium was replaced with fresh complete growth medium. The transfected cells were collected three days post-transfection for qRT-PCR or flow cytometry analysis. Sequences of tigRNAs for gene activation are listed in **Supplementary Table S3**.

### Flow cytometry

To analyze fluorescent protein expression, cells were dissociated using 0.05% Trypsin-EDTA (Gibco, USA), resuspended in DMEM with 10% FBS, and analyzed on an Attune NxT flow cytometer (Thermo Fisher Scientific, USA). Gating strategies for all the samples are shown in **Supplementary Fig. 7**.

### Quantitative real-time polymerase chain reaction (qRT-PCR)

Total RNA was isolated using Trizol reagent (Vazyme, China) and 500 ng of RNA was used as the template for cDNA synthesis (Vazyme, China). The reaction reagent was transferred using Echo 525 Liquid Handler (Beckman Coulter, USA). qRT-PCR was performed on the Applied Biosystems QuantStudio™ 7 Pro (Thermo Fisher Scientific, USA) using Taq Pro Universal SYBR qPCR Master Mix (Vazyme, China). qRT-PCR amplifications were performed in triplicates for each sample. The relative mRNA expression level was calculated using the 2^-ΔΔCt^ method. The housekeeping gene *glyceraldehyde phosphate dehydrogenase* (*GAPDH*) was used as an internal control. Data were represented as fold change to the dTasR control group. All qPCR primers are listed in **Supplementary Table S4**.

### Yeast culture, transformation, and mCherry measurement

The *S. cerevisiae* strain CT (CEN.PK2-1c-ura3::URA3-CYC1p-mCherry-TEF1t-TEF1p-mVenus-PGK1t, a kind gift from Dr. Jiazhang Lian, Zhejiang University) was used as the reporter strain for mCherry activation experiments. The strain was cultivated in YPD medium (10 g/liter yeast extract, 20 g/liter tryptone, 20 g/liter glucose) before transformation. Plasmid transformation of CT (1 μg of plasmid per transformation) was carried out using the LiAc/SS carrier DNA/PEG method. After transformation, cells were incubated in SC-L medium for 4 days and then inoculated into fresh SC-Leu medium with an initial OD of 0.2 and cultivated for 2 days at 30 °C, 250 rpm. The transformed cells were then collected, resuspended in PBS, and analyzed on Attune NxT flow cytometer (Thermo Fisher Scientific, USA).

### Transcriptome profiling by RNA sequencing

HEK293T cells were transfected with dTasR-10×GCN4, scFv-P65-HSF1 and 4 pooled tigRNA plasmids targeting the *HBG* locus and collected for RNA extraction three days post-transfection. Total RNA was isolated with Trizol Reagent. mRNA was enriched using oligo T (dT) beads and fragmented for library construction. The constructed sequencing libraries were sequenced on the Illumina NovaSeq platform with paired-end 150 bp read (GENEWIZ, China). Adapters and low-quality reads were trimmed using Fastp (v1.1.0). The paired-end clean reads were aligned to human transcriptomes (Gencode release 48) and quantified using Salmon (1.10.3). Genes were filtered to include only those with counts > 0. Transcripts per kilobase million (TPM) was calculated based on the length of the gene exon to quantify gene expression levels.

### Statistics

Statistical analyses were carried out with GraphPad Prism software (version 10). Error bars represent the standard error of the mean (S.E.M.) and results were presented as mean ± S.E.M.

## Code availability

All computational tools used for analyses of the RNA-seq data are available from provided references in Methods. Custom batch scripts used for execution of these computational tools can be found in **Supplementary Code**.

## Data availability

The raw reads of the RNA-seq data were deposited into the Sequence Read Archive (SRA) database with accession number PRJNA1452194 at the National Center for Biotechnology Information (NCBI).

