## Supplementary Information for "Repurposing the TIGR-Tas system for programmable transcription activation"

### SUPPLEMENTARY FIGURES

```
TaTasR      8 LAVDWSHEERKLAIFDGKKIRKKLPEPSSDVIIVAENIPQKYAAPFIEVGAKVLRCSTNATADARKNY-QKKVDAAFAKN 86
              +A+DW+  E KLA++DGKK+ K +P+  +   I AENIP K+AA +++ G  + RC  N TA  RK   Q+K DA
ParTasR     4 VAIDWAFAEEKLAVYDGKKVLKAVPKLKAGDEIFAENIPMKHAAQWLKDGIIIRRCRPNDTAALRKEIGQEKTDAL---- 79

TaTasR     87 DENDSKVIWALYQTHPELFREMKLEPPLSSYYAIFKDYQEVRIRTGNRLYSDRTDAMEEFFKIVKKGEHELKKAVDKELE 166
                 D+K+IW L + HPE FRE K +P L++ Y  FK+ Q  R+   NR+++   +        ++K E ++ KA++KEL+
ParTasR    79 ---DAKLIWQLAEAHPEKFREWKGDPQLTTLYRAFKEVQRCRVGQSNRVWAKGEETATAVLDDLEKTERKIVKAIEKELK 156

TaTasR    167 NHPVYTQWLQHIKGIGPVVAGGLISLIG--DIDRFDSVSKLWAYAGYSVDNGKVQKRKKGVASNWKNKIRTHCYNIV-DS 243
              ++ V+  WL  IKGIG    GGL+ LI    I+    VS LW   G  V  GK  +R KG   ++  + +T    ++ D
ParTasR   157 SYKVW-DWLSQIKGIGVATGGGLVGLIAKYGIENIRQVSSLWHLFGLHVVEGKAPRRTKGEEVSYVPEAKTLVLGVIADC 235

TaTasR    244 FIKQRTSVYRELYDAEKARQ----RPKVE----------SD-----GHAHNRAVRKVAKVFLQHYWVVSRELAGFSVSKP 304
              FIKQR+ VYR++YD EKARQ     P+ E          SD      HAH RA+RK+ K+F+QH W+  R   G
ParTasR   236 FIKQRSPVYRDIYDQEKARQLEIEYPEGELASKFLGYKKSDTHLSLNHAHRRAIRKMMKIFVQHVWLAWRTCEGLETRPL 315

TaTasR    305 WILEHGGHVDYIKPPHWNKVEIKP 328
              +  E+ GH  +I+PP    V+I+P
ParTasR   316 YCHEYLGHEHFIEPP----VKIQP 335
```

**Figure S1. Sequence alignment of TaTasR and ParTasR.** "+" indicates biochemically similar amino acid residues. The mutated residue D for nuclease inactivation is denoted with a red box.

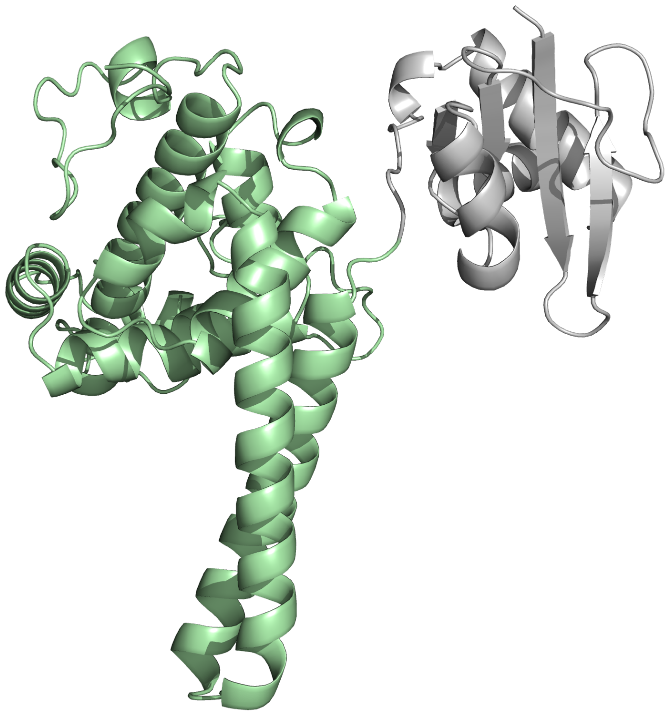

**Figure S2. Alphfold3 predicted structure of ParTasR monomer.** Gray, RuvC domain deleted in tTasR; green, the coiled-coil domain and the tigRNA-binding domain.

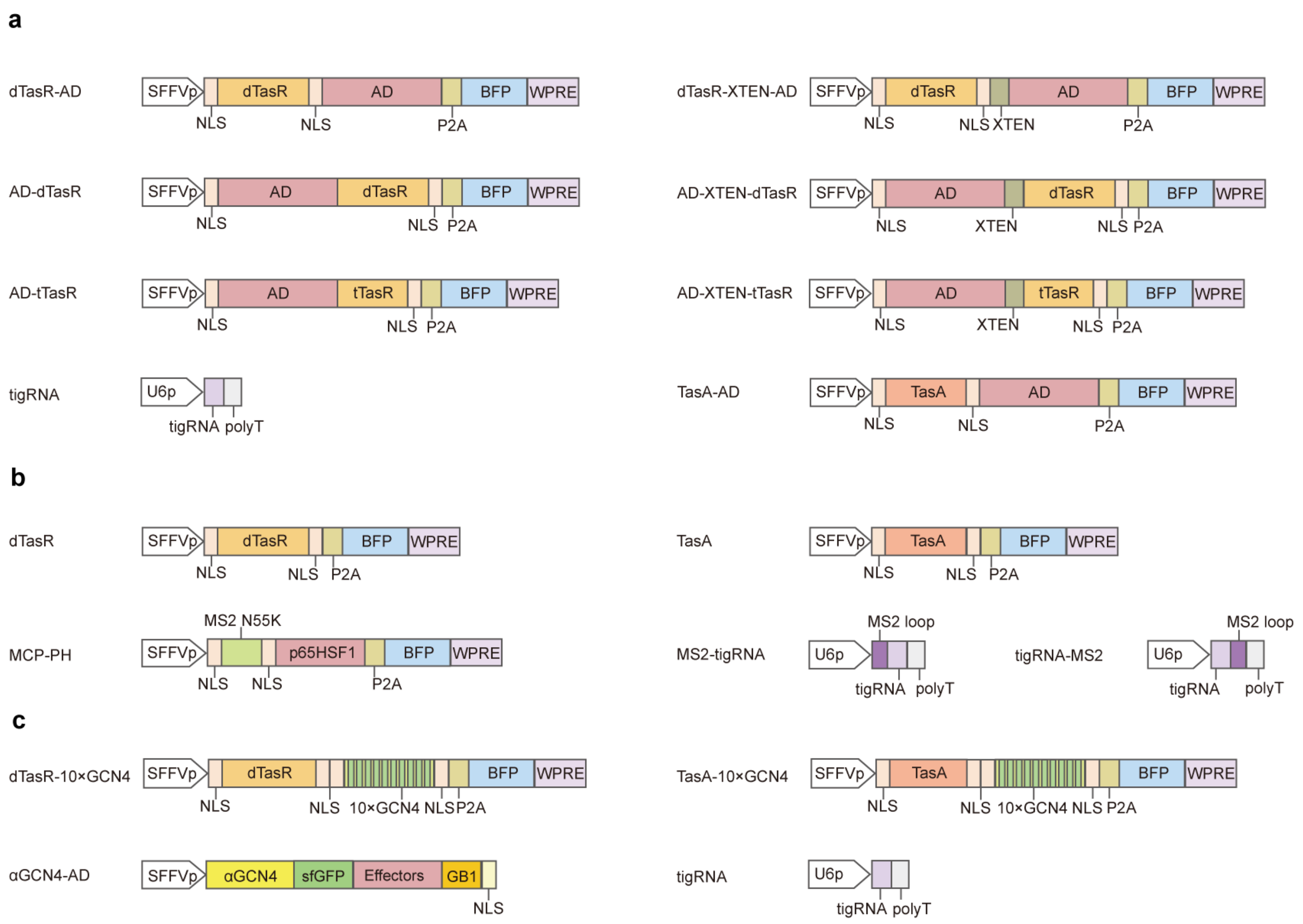


**Figure S3. Diagram of constructed transcription activators based on TasR and TasA in this study.** **a**, activators based on TasR and TasA direction fusions. **b**, MS2-tigRNA and tigRNA-MS2. **c**, SunTag systems based on TasA and TasR.

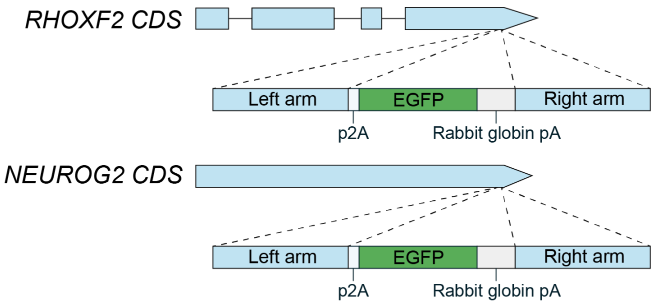


**Figure S4. Schematic of EGFP-tagged endogenous reporter cell lines.** p2A-EGFP fragment was knocked in by CRISPR-Cas9 mediated homology directed repair. CDS, coding sequence.

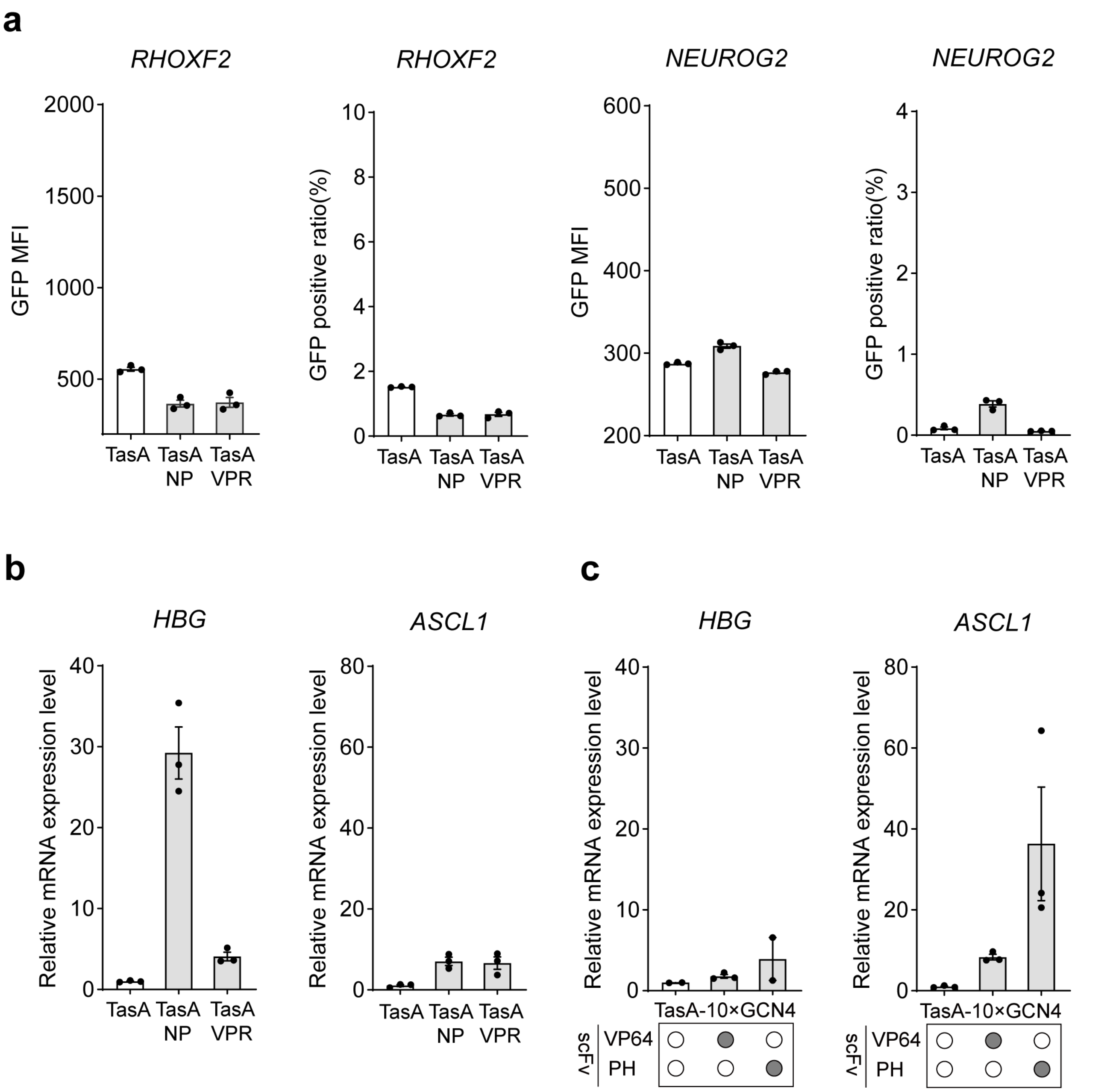


**Figure S5. Activation effects of TasA-AD direct fusions and TasA-SunTag at EGFP reporter loci and endogenous genes. a**, the activation effects of TasA-AD fusions at *RHOXF2-EGFP* reporter locus and *NEUROG2-EGFP* reporter locus. The data were graphed as mean ± S.E.M and represent three biological repeats. **b**, relative expression levels of different TasA-AD fusions at *HBG* and *ASCL1* in HEK293T. The data were graphed as mean ± S.E.M and represent three biological replicates. **c**, relative expression levels of different TasA-SunTag fusions at *HBG* and *ASCL1* in HEK293T. The data were graphed as mean ± S.E.M and represent at least two biological replicates. The housekeeping gene *glyceraldehyde phosphate dehydrogenase* (GAPDH) was used as an internal control for the normalization of qRT-PCR data.

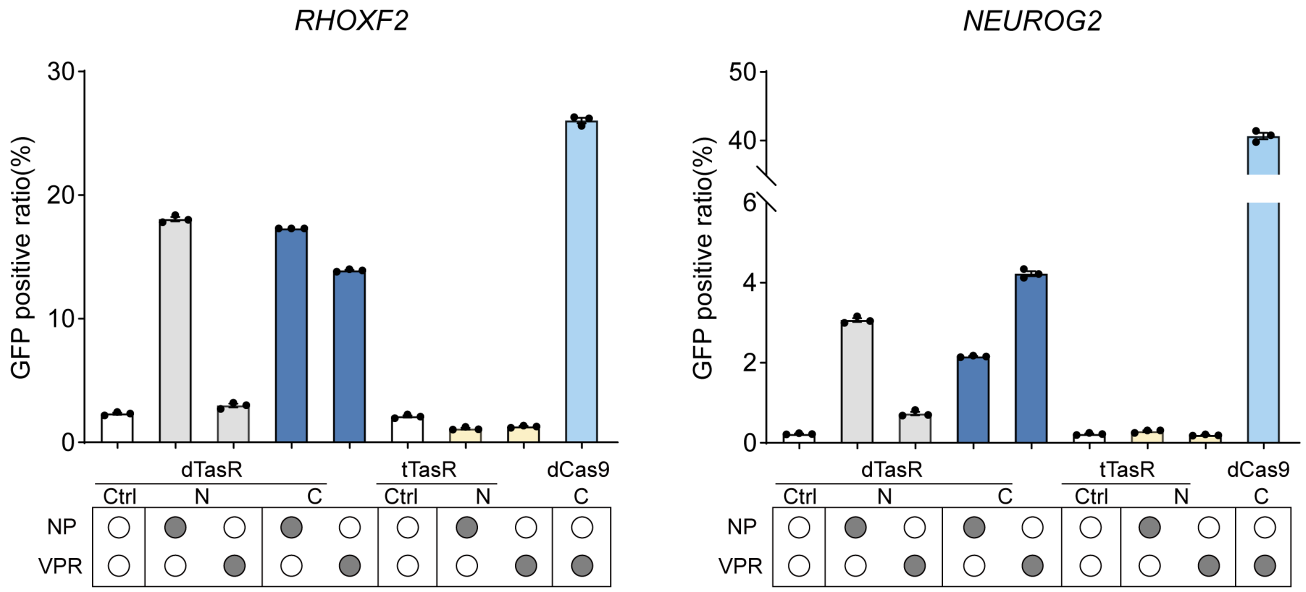


**Figure S6. The EGFP positive ratio with different dTasR and tTasR fusions at *RHOXF2-EGFP* and *NEUROG2-EGFP* reporter loci.** Ctrl, dTasR or tTasR without ADs. The data were graphed as mean ± S.E.M and represent three biological replicates.

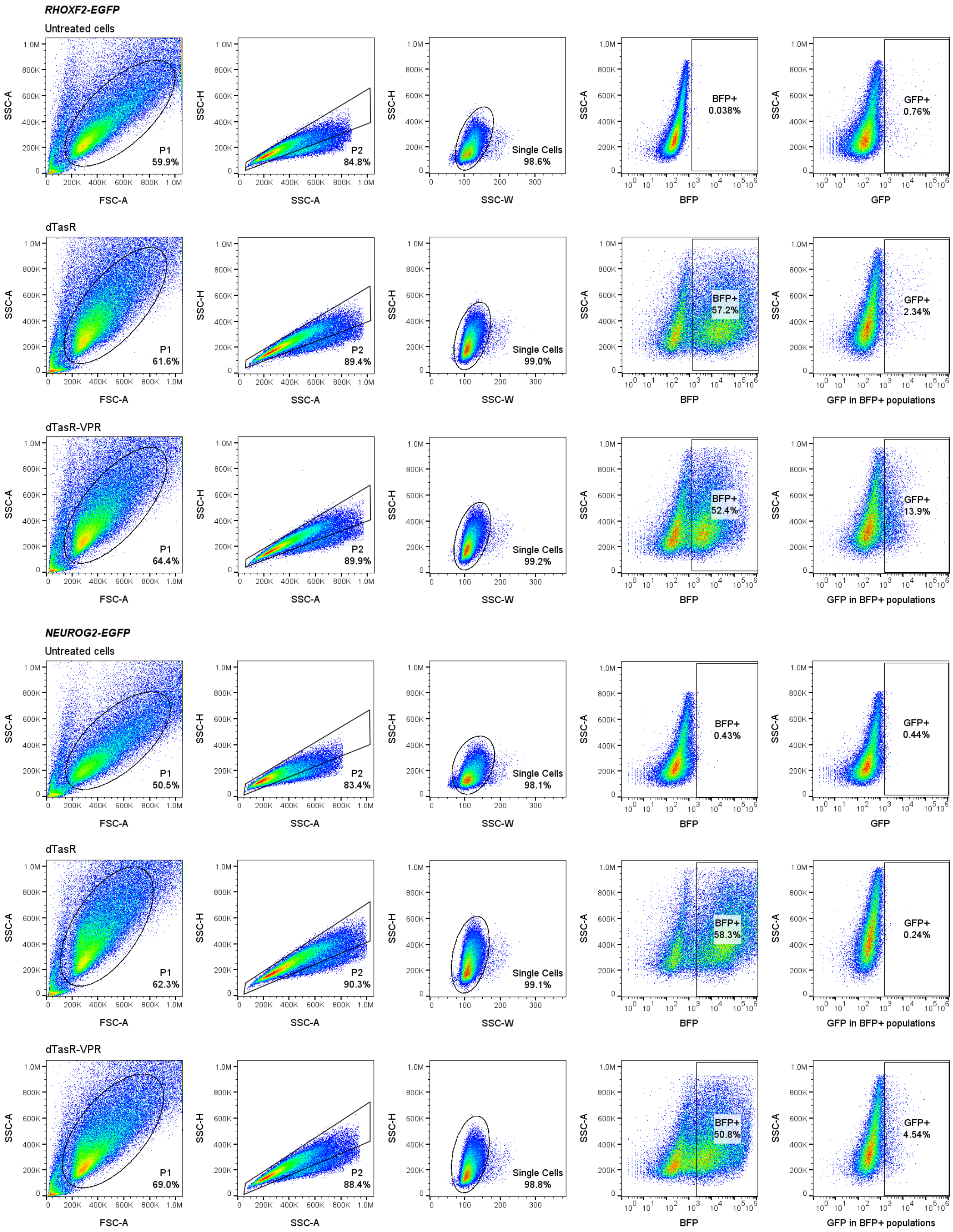


**Figure S7. Flow cytometry gating strategy used for assessing transcription activation in EGFP reporter cell lines.** Untreated cells were used to gate BFP and GFP positive populations.

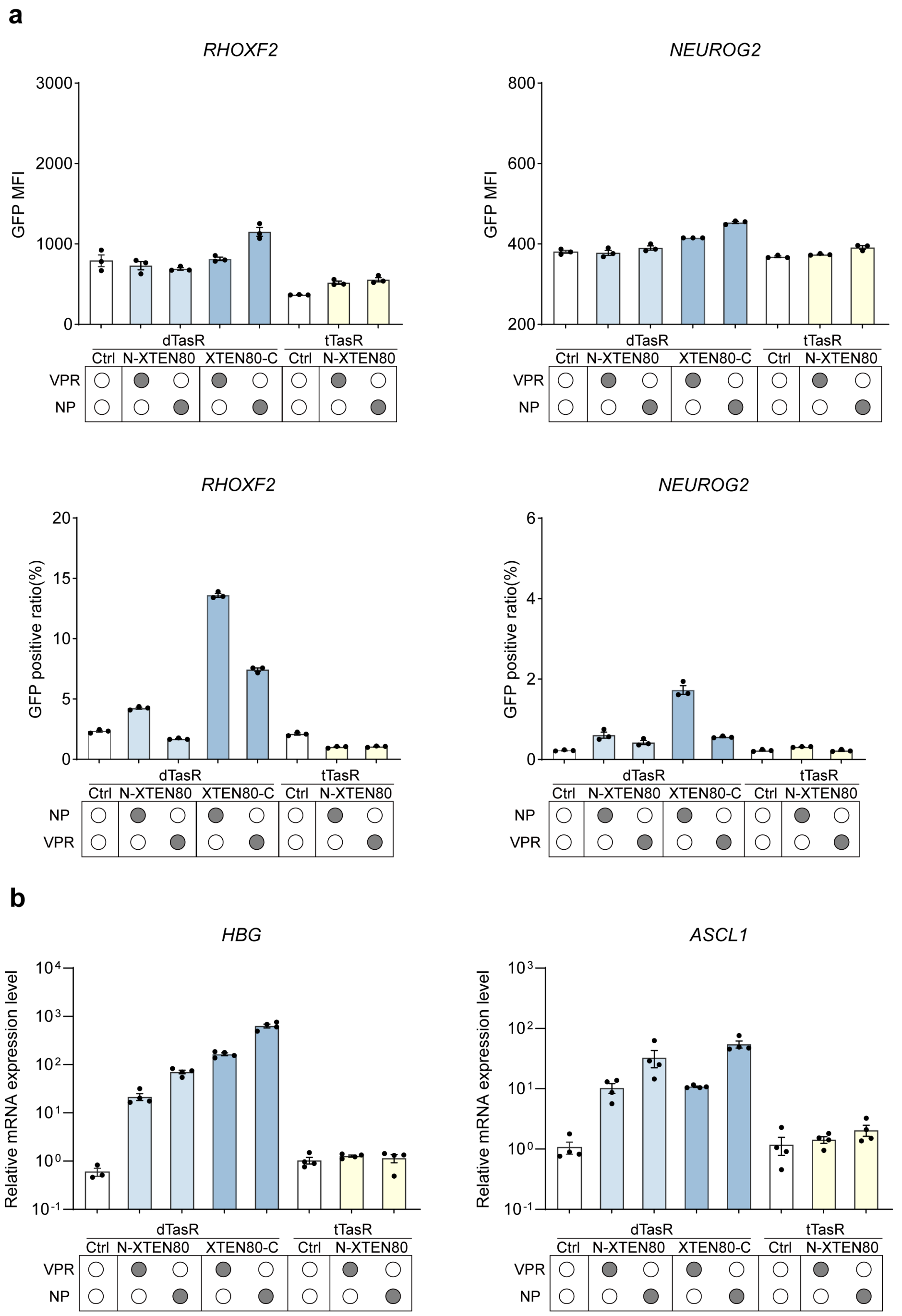


**Figure S8. The effects of XTEN80 linkers on dTasR and tTasR direct AD fusions at two EGFP reporter loci and endogenous genes. a**, effects at *RHOXF2*-EGFP and *NEUROG2*-EGFP reporter loci. The data were graphed as mean ± S.E.M and represent three biological replicates. MFI, mean fluorescence intensity. **b**, relative expression levels at *HBG* and *ASCL1* in HEK293T. The housekeeping gene *glyceraldehyde phosphate dehydrogenase* (GAPDH) was used as an internal control for the normalization of qRT-PCR data. The data were graphed as mean ± S.E.M and represent four biological replicates.

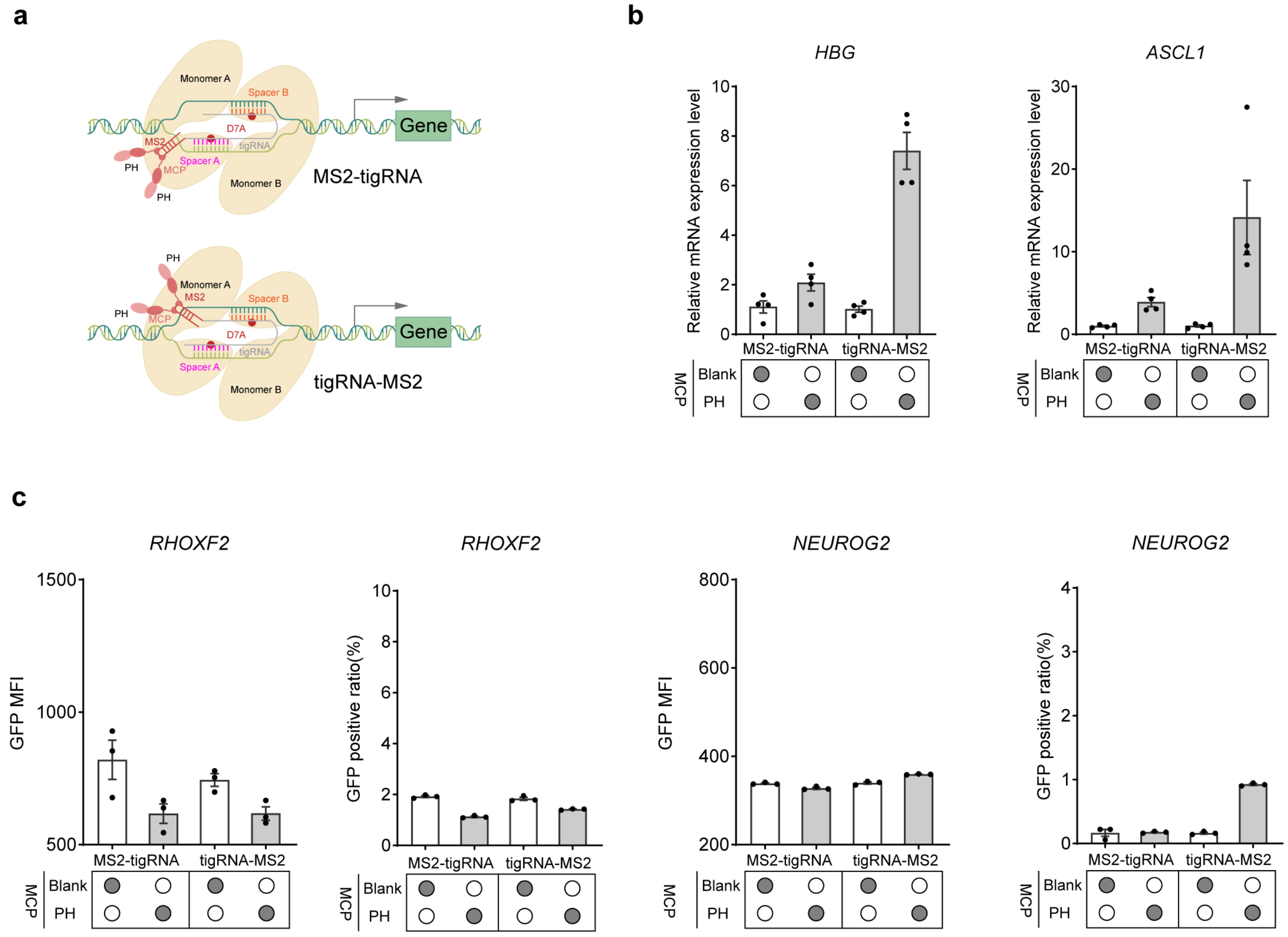


**Figure S9. Activation effects of MS2-tigRNA and tigRNA-MS2 at two EGFP reporter loci and endogenous genes.** **a**, schematic of transcription activators with the fusion of the MS2 aptamer to either end of the tigRNA of TasR. **b**, relative expression levels of MS2-tigRNA and tigRNA-MS2 at *HBG* and *ASCL1* in HEK293T. The housekeeping gene *glyceraldehyde phosphate dehydrogenase* (GAPDH) was used as an internal control for the normalization of qRT-PCR data. The data were graphed as mean ± S.E.M and represent four biological replicates. **c**, effects of MS2-tigRNA and tigRNA-MS2 at *RHOXF2*-EGFP and *NEUROG2*-EGFP reporter loci. The data were graphed as mean ± S.E.M and represent three biological repeats. MFI, mean fluorescence intensity.

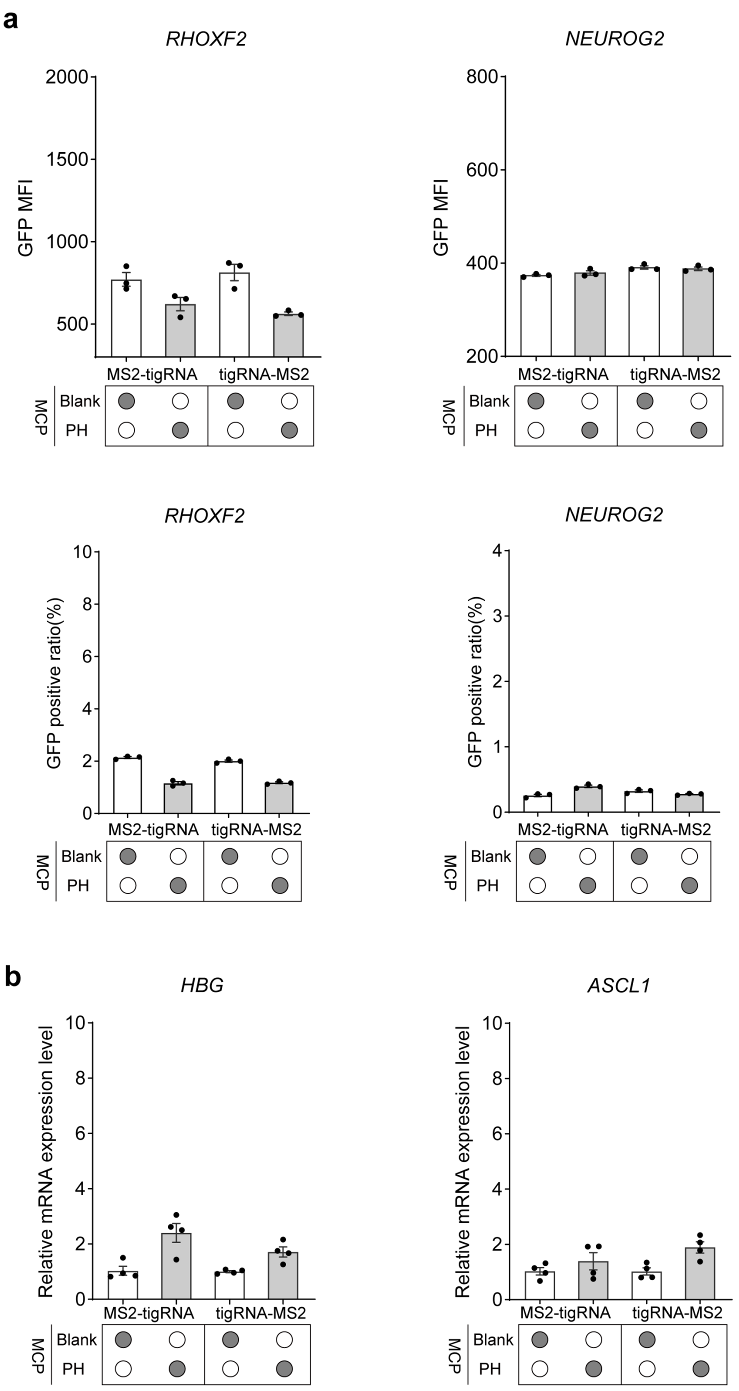


**Figure S10. Activation effects of MS2-tigRNA and tigRNA-MS2 based on TasA at two EGFP reporter loci and endogenous genes. a**, effects at *RHOXF2-EGFP* and *NEUROG2-EGFP* reporter loci. The data were graphed as mean ± S.E.M and represent three biological replicates. MFI, mean fluorescence intensity. **b**, relative expression levels at *HBG* and *ASCL1* in HEK293T. The data were graphed as mean ± S.E.M and represent four biological replicates. The housekeeping gene *glyceraldehyde phosphate dehydrogenase* (GAPDH) was used as an internal control for the normalization of qRT-PCR data.

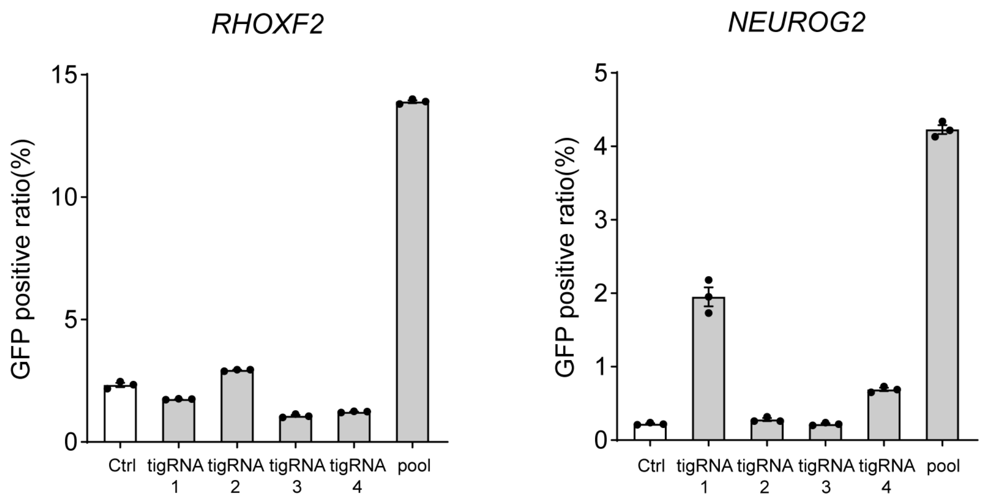


**Figure S11. The EGFP positive ratio of dTasR-VPR using either individual or pooled tigRNAs at *RHOXF2-EGFP* and *NEUROG2-EGFP* reporter loci.** Ctrl, dTasR without ADs. The data were graphed as mean ± S.E.M and represent three biological replicates.

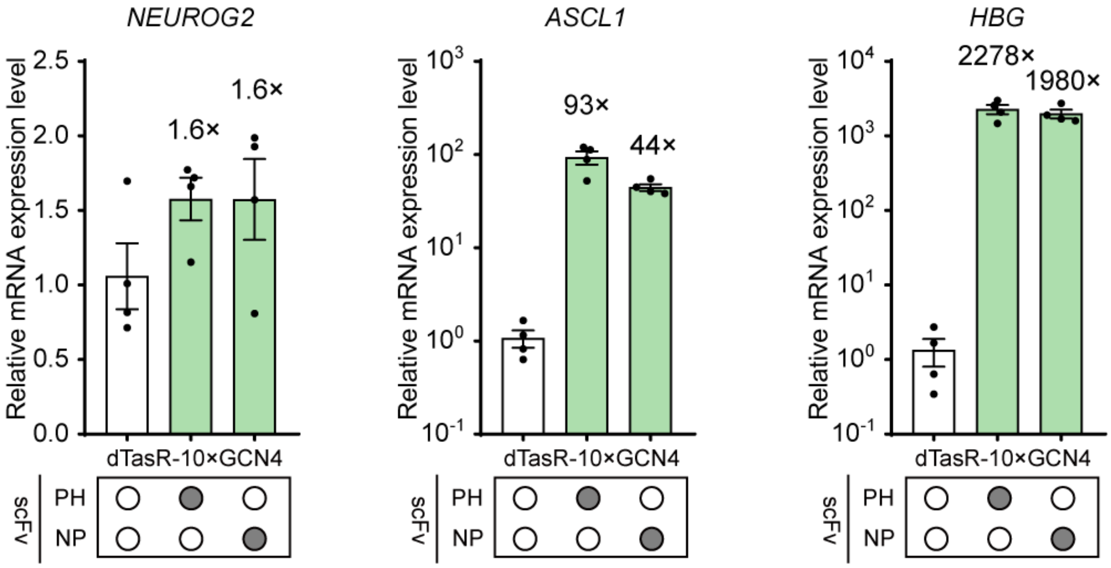


**Figure S12. Multiplex activation of three endogenous genes *NEUROG2*, *ASCL1* and *HBG* with dTasR-SunTag using pooled tigRNAs**. Ctrl, dTasR-SunTag without AD. The housekeeping gene *glyceraldehyde phosphate dehydrogenase* (GAPDH) was used as an internal control for the normalization of qRT-PCR data. The data were graphed as mean ± S.E.M and represent four biological replicates.

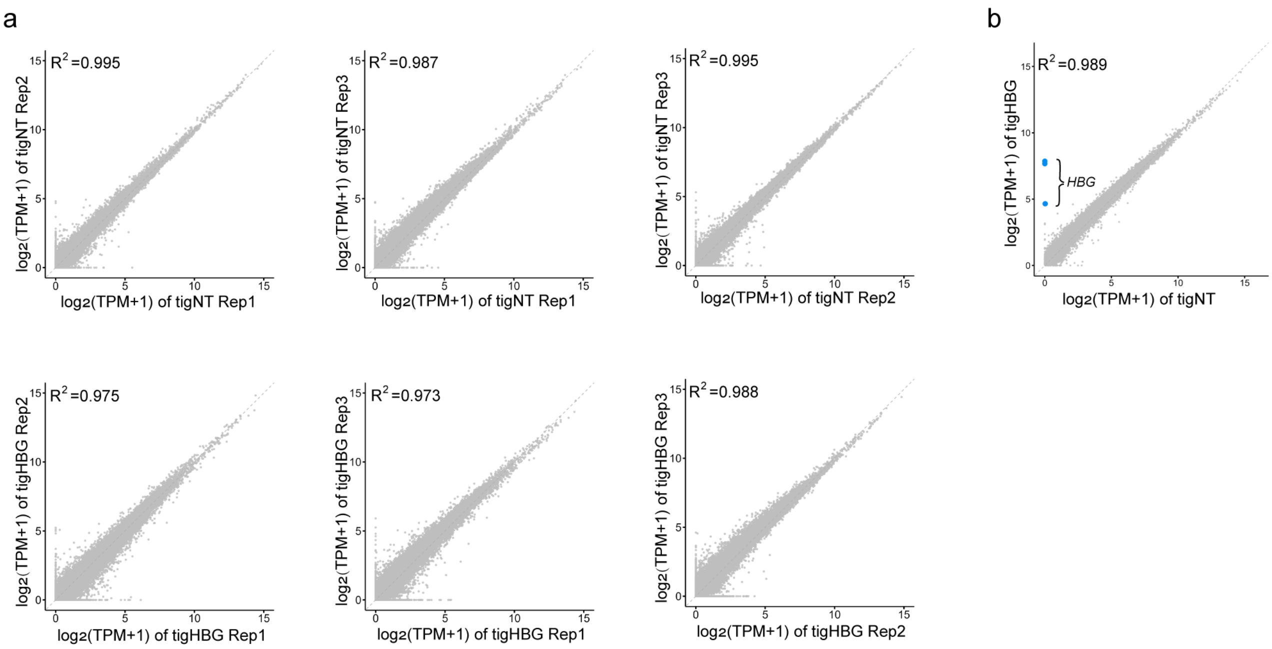


**Figure S13. RNA-seq profiling of dTasR-SunTag in HEK293T cells. a**, Scatter plots comparing $\log_2$(TPM+1) values for biological replicates for each condition were shown. First row: dTasR-SunTag-PH plus non-targeting tigRNAs. Second row: dTasR-SunTag-PH plus *HBG*-targeting tigRNAs. The calculated Pearson correlation coefficient for each condition is at top left. **b**, Scatterplot comparing $\log_2$(TPM+1) values of dTasR-SunTag-PH with targeting tigRNAs versus non-targeting tigRNAs. TPM, transcripts per kilobase million.

### SUPPLEMENTARY TABLES

**Table S1. The amino acid sequences of activators constructed in this study.**

| |
|---|
| **dTasR-NP:** SV40 Nuclear Localization Signal, *Parcubacteria* dTasR (D7A), NP |
| PKKKRKVGIHGVPGGLEGGGGSGGTASMRQVAIAWAFAEEKLAVYDGKKVLKAVPKLKA<br>GDEIFAENIPMKHAAQWLKDGIIIRRCRPNDTAALRKEIGQEKTDALDAKLIWQLAEAHPE<br>KFREWKGDPQLTTLYRAFKEVQRCRVGQSNRVWAKGEETATAVLDDLEKTERKIVKAIEK<br>ELKSYKVWDWLSQIKGIGVATGGGLVGLIAKYGIENIRQVSSLWHLFGLHVVEGKAPRRT<br>KGEEVSYVPEAKTLVLGVIADCFIKQRSPVYRDIYDQEKARQLEIEYPEGELASKFLGYKK<br>SDTHLSLNHAHRRAIRKMMKIFVQHVWLAWRTCEGLETRPLYCHEYLGHEHFIEPPVKIQ<br>PEGSSPKKKRKVGSEGQSDERALLDQLHTLLSNTDATGLEEIDRALGIPELVNQGQALEPK<br>QGSGSGSHEKFPSDLDLDMFNGSLECDMESIIRSELMDADGLDFNFDSGSGSGSFVTLKDV<br>GMDFTLGDWEQLGLEQGDTFWDTALDNCQDLFLLGGGGSPSGQISNQALALAPSSAPVL<br>AQTMVPSSAMVPLAQPPAPAPVLTPGPPQSLSAPVPKSTQAGEGTLSEALLHLQFDADEDL<br>GALLGNSTDPGVFTDLASVDNSEFQQLLNQGVSMSHSTAEPMLMEYPEAITRLVTGSQRP<br>PDPAPTPLGTSGLPNGLSGDEDFSSIADMDFSALLSQISSSGQGGGGSGFSVDTSALLDLFSP<br>SVTVPDMSLPDLDSSLASIQELLSPQEPPRPPEAENSSPDSGKQLVHYTAQPLFLLDPGSVD<br>TGSNDLPVLFELGEGSYFSEGDGFAEDPTISLLTGSEPPKAKDPTVS |
| **dTasR-VPR:** SV40 Nuclear Localization Signal, *Parcubacteria* dTasR (D7A), VPR |
| PKKKRKVGIHGVPGGLEGGGGSGGTASMRQVAIAWAFAEEKLAVYDGKKVLKAVPKLKA<br>GDEIFAENIPMKHAAQWLKDGIIIRRCRPNDTAALRKEIGQEKTDALDAKLIWQLAEAHPE<br>KFREWKGDPQLTTLYRAFKEVQRCRVGQSNRVWAKGEETATAVLDDLEKTERKIVKAIEK<br>ELKSYKVWDWLSQIKGIGVATGGGLVGLIAKYGIENIRQVSSLWHLFGLHVVEGKAPRRT<br>KGEEVSYVPEAKTLVLGVIADCFIKQRSPVYRDIYDQEKARQLEIEYPEGELASKFLGYKK<br>SDTHLSLNHAHRRAIRKMMKIFVQHVWLAWRTCEGLETRPLYCHEYLGHEHFIEPPVKIQ<br>PEGSSPKKKRKVGSEASGSGRADALDDFDLDMLGSDALDDFDLDMLGSDALDDFDLDM<br>LGSDALDDFDLDMLINSRSSGSPKKKRKVGSQYLPDTDDRHRIEEKRKRTYETFKSIMKKS<br>PFSGPTDPRPPPRRIAVPSRSSASVPKPAPQPYPFTSSLSTINYDEFPTMVFPSGQISQASALA<br>PAPPQVLPQAPAPAPAPAMVSALAQAPAPVPVLAPGPPQAVAPPAPKPTQAGEGTLSEALLQ<br>LQFDDEDLGALLGNSTDPAVFTDLASVDNSEFQQLLNQGIPVAPHTTEPMLMEYPEAITRL<br>VTGAQRPPDPAPAPLGAPGLPNGLLSGDEDFSSIADMDFSALLGSGSGSRDSREGMFLPKP<br>EAGSAISDVFEGREVCQPKRIRPFHPPGSPWANRPLPASLAPTPTGPVHEPVGSLTPAPVPQP<br>LDPAPAVTPEASHLLEDPDEETSQAVKALREMADTVIPQKEEAAICGQMDLSHPPPRGHLD<br>ELTTTLESMTEDLNLDSPLTPELNEILDTFLNDECLLHAMHISTGLSIFDTSLF |
| **dTasR-10×GCN4:** SV40 Nuclear Localization Signal, *Parcubacteria* dTasR (D7A), 10×GCN4-GS linker |
| PKKKRKVGIHGVPGGLEGGGGSGGTASMRQVAIAWAFAEEKLAVYDGKKVLKAVPKLKA<br>GDEIFAENIPMKHAAQWLKDGIIIRRCRPNDTAALRKEIGQEKTDALDAKLIWQLAEAHPE<br>KFREWKGDPQLTTLYRAFKEVQRCRVGQSNRVWAKGEETATAVLDDLEKTERKIVKAIEK |

ELKSYKVWDWLSQIKGIGVATGGGLVGLIAKYGIENIRQVSSLWHLFGLHVVEGKAPRRT
KGEEVSYVPEAKTLVLGVIADCFIKQRSPVYRDIYDQEKARQLEIEYPEGELASKFLGYKK
SDTHLSLNHAHRRAIRKMMKIFVQHVWLAWRTCEGLETRPLYCHEYLGHEHFIEPPVKIQ
PEGSSPKKKRKVGSPKKKRKVNGPTDAAEEELLSKNYHLENEVARLKKGSGSGEELLSKN
YHLENEVARLKKGSGSGEELLSKNYHLENEVARLKKGSGSGEELLSKNYHLENEVARLKK
GSGSGEELLSKNYHLENEVARLKKGSGSGEELLSKNYHLENEVARLKKGSGSGEELLSKN
YHLENEVARLKKGSGSGEELLSKNYHLENEVARLKKGSGSGEELLSKNYHLENEVARLKK
GSGSGEELLSKNYHLENEVARLKKGSGSGTAVNIGGGTGPMDLQPKKKRKV

**scFv-VP64:** scFv, super-folder GFP, VP64, B1 domain of Streptococcal protein G, HTLV-1 Rex Nuclear Localization Signal

MGPDIVMTQSPSSLSASVGDRVTITCRSSTGAVTTSNYASWVQEKPGKLFKGLIGGTNNRA
PGVPSRFSGSLIGDKATLTISSLQPEDFATYFCALWYSNHWVFGQGTKVELKRGGGGSGGG
GSGGGGSSGGGSEVKLLESGGGLVQPGGSLKLSCAVSGFSLTDYGVNWVRQAPGRGLEW
IGVIWGDGITDYNSALKDRFIISKDNGKNTVYLQMSKVRSDDTALYYCVTGLFDYWGQGT
LVTVSSYPYDVPDYAGGGGGSGGGGSGGGGSGGGGSLDPGGGGSGSKGEELFTGVVPILV
ELDGDVNGHKFSVRGEGEGDATNGKLTLKFICTTGKLPVPWPTLVTTLTYGVQCFSRYPD
HMKRHDFFKSAMPEGYVQERTISFKDDGTYKTRAEVKFEGDTLVNRIELKGIDFKEDGNI
LGHKLEYNFNSHNVYITADKQKNGIKANFKIRHNVEDGSVQLADHYQQNTPIGDGPVLLP
DNHYLSTQSVLSKDPNEKRDHMVLLEFVTAAGITHGMDELYKGGGRTGGGGGGEFDALD
DFDLDMLGSDALDDFDLDMLGSDALDDFDLDMLGSDALDDFDLDMLEFGSGGGSRTEE
YKLILNGKTLKGETTTEAVDAATAEKVFKQYANDNGVDGEWTYDDATKTFTVTEGGGSG
GGTSPKTRRRPRRSQRKRPPT

**scFv-P65-HSF1:** scFv, super-folder GFP, P65-HSF1, B1 domain of Streptococcal protein G, HTLV-1 Rex Nuclear Localization Signal

MGPDIVMTQSPSSLSASVGDRVTITCRSSTGAVTTSNYASWVQEKPGKLFKGLIGGTNNRA
PGVPSRFSGSLIGDKATLTISSLQPEDFATYFCALWYSNHWVFGQGTKVELKRGGGGSGGG
GSGGGGSSGGGSEVKLLESGGGLVQPGGSLKLSCAVSGFSLTDYGVNWVRQAPGRGLEW
IGVIWGDGITDYNSALKDRFIISKDNGKNTVYLQMSKVRSDDTALYYCVTGLFDYWGQGT
LVTVSSYPYDVPDYAGGGGGSGGGGSGGGGSGGGGSLDPGGGGSGSKGEELFTGVVPILV
ELDGDVNGHKFSVRGEGEGDATNGKLTLKFICTTGKLPVPWPTLVTTLTYGVQCFSRYPD
HMKRHDFFKSAMPEGYVQERTISFKDDGTYKTRAEVKFEGDTLVNRIELKGIDFKEDGNI
LGHKLEYNFNSHNVYITADKQKNGIKANFKIRHNVEDGSVQLADHYQQNTPIGDGPVLLP
DNHYLSTQSVLSKDPNEKRDHMVLLEFVTAAGITHGMDELYKGGGRTGGGGGGEFPSGQ
ISNQALALAPSSAPVLAQTMVPSSAMVPLAQPPAPAPVLTPGPPQSLSAPVPKSTQAGEGTL
SEALLHLQFDADEDLGALLGNSTDPGVFTDLASVDNSEFQQLLNQGVSMSHSTAEPMLM
EYPEAITRLVTGSQRPPDPAPTPLGTSGLPNGLSGDEDFSSIADMDFSALLSQISSSGQGGGG
SGFSVDTSALLDLFSPSVTVPDMSLPDLDSSLASIQELLSPQEPPRPPEAENSSPDSGKQLVH
YTAQPLFLLDPGSVDTGSNDLPVLFELGEGSYFSEGDGFAEDPTISLLTGSEPPKAKDPTVS
EFGSGGGSRTEEYKLILNGKTLKGETTTEAVDAATAEKVFKQYANDNGVDGEWTYDDAT
KTFTVTEGGGSGGGTSPKTRRRPRRSQRKRPPT

**scFv-NP:** scFv, super-folder GFP, NP, B1 domain of Streptococcal protein G, HTLV-1 Rex NLS

MGPDIVMTQSPSSLSASVGDRVTITCRSSTGAVTTSNYASWVQEKPGKLFKGLIGGTNNRA
PGVPSRFSGSLIGDKATLTISSLQPEDFATYFCALWYSNHWVFGQGTKVELKRGGGGSGGG
GSGGGGSSGGGSEVKLLESGGGLVQPGGSLKLSCAVSGFSLTDYGVNWVRQAPGRGLEW
IGVIWGDGITDYNSALKDRFIISKDNGKNTVYLQMSKVRSDDTALYYCVTGLFDYWGQGT
LVTVSSYPYDVPDYAGGGGGSGGGGSGGGGSGGGGSLDPGGGGSGSKGEELFTGVVPILV
ELDGDVNGHKFSVRGEGEGDATNGKLTLKFICTTGKLPVPWPTLVTTLTYGVQCFSRYPD
HMKRHDFFKSAMPEGYVQERTISFKDDGTYKTRAEVKFEGDTLVNRIELKGIDFKEDGNI
LGHKLEYNFNSHNVYITADKQKNGIKANFKIRHNVEDGSVQLADHYQQNTPIGDGPVLLP
DNHYLSTQSVLSKDPNEKRDHMVLLEFVTAAGITHGMDELYKGGGRTGGGGGGEFEGQS
DERALLDQLHTLLSNTDATGLEEIDRALGIPELVNQGQALEPKQGSGSGSHEKFPSDLDLD
MFNGSLECDMESIIRSELMDADGLDFNFDSGSGSGSFVTLKDVGMDFTLGDWEQLGLEQ
GDTFWDTALDNCQDLFLLGGGGSPSGQISNQALALAPSSAPVLAQTMVPSSAMVPLAQPP
APAPVLTPGPPQSLSAPVPKSTQAGEGTLSEALLHLQFDADEDLGALLGNSTDPGVFTDLA
SVDNSEFQQLLNQGVSMSHSTAEPMLMEYPEAITRLVTGSQRPPDPAPTPLGTSGLPNGLS
GDEDFSSIADMDFSALLSQISSSGQGGGGSGFSVDTSALLDLFSPSVTVPDMSLPDLDSSLA
SIQELLSPQEPPRPPEAENSSPDSGKQLVHYTAQPLFLLDPGSVDTGSNDLPVLFELGEGSY
FSEGDGFAEDPTISLLTGSEPPKAKDPTVSEFGSGGGSRTEEYKLILNGKTLKGETTTEAVD
AATAEKVFKQYANDNGVDGEWTYDDATKTFTVTEGGGSGGGTSPKTRRRPRRSQRKRPP
T

**MCP-P65-HSF1:** SV40 Nuclear Localization Signal, MS2 bacteriophage coat protein, P65-HSF1

PKKKRKVGIHGVPGGLEGGGGSGGTASASNFTQFVLVDNGGTGDVTVAPSNFANGVAEWI
SSNSRSQAYKVTCSVRQSSAQKRKYTIKVEVPKVATQTVGGVELPVAAWRSYLNMELTIPI
FATNSDCELIVKAMQGLLKDGNPIPSAIAANSGIYSPKKKRKVGSPSGQISNQALALAPSSA
PVLAQTMVPSSAMVPLAQPPAPAPVLTPGPPQSLSAPVPKSTQAGEGTLSEALLHLQFDAD
EDLGALLGNSTDPGVFTDLASVDNSEFQQLLNQGVSMSHSTAEPMLMEYPEAITRLVTGS
QRPPDPAPTPLGTSGLPNGLSGDEDFSSIADMDFSALLSQISSSGQGGGGSGFSVDTSALLD
LFSPSVTVPDMSLPDLDSSLASIQELLSPQEPPRPPEAENSSPDSGKQLVHYTAQPLFLLDPG
SVDTGSNDLPVLFELGEGSYFSEGDGFAEDPTISLLTGSEPPKAKDPTVS

**TasA-NP:** SV40 Nuclear Localization Signal, *Flavonifractor plautii* TasA, NP

PKKKRKVGIHGVPGGLEGGGGSGGTASMDIQKNRIRNIVGGIYDIQKLRIATGNRIVASLRP
GLVEEVKEGEEDTKYLPAILSEYRRITDYFVSEFEGRGSIEKAITPNNPEYIKSRLDYDLVTS
YKRLLETEEGLTKVAEREVKAHPMWDAFFAGVKGCGPLMSAVCLAYFDPYKARHASSFW
RYAGLDVQRDPDKDKMRGVGKWYTEERPYIDKDGKEQMKKSLTYNPFLKTKLVGVLGS
AFLRAKDSYYGKVYYDYKNRLDNRDEDFSPIVKHRMATRYAVKMFLRDMWVVWRELE
GLEVTEPYEVAKLGHKPHHSSSPKKKRKVGSEGQSDERALLDQLHTLLSNTDATGLEEIDR
ALGIPELVNQGQALEPKQGSGSGSHEKFPSDLDLDMFNGSLECDMESIIRSELMDADGLDF
NFDSGSGSGSFVTLKDVGMDFTLGDWEQLGLEQGDTFWDTALDNCQDLFLLGGGGSPSG

QISNQALALAPSSAPVLAQTMVPSSAMVPLAQPPAPAPVLTPGPPQSLSAPVPKSTQAGEG
TLSEALLHLQFDADEDLGALLGNSTDPGVFTDLASVDNSEFQQLLNQGVSMSHSTAEPML
MEYPEAITRLVTGSQRPPDPAPTPLGTSGLPNGLSGDEDFSSIADMDFSALLSQISSSGQGG
GGSGFSVDTSALLDLFSPSVTVPDMSLPDLDSSLASIQELLSPQEPPRPPEAENSSPDSGKQL
VHYTAQPLFLLDPGSVDTGSNDLPVLFELGEGSYFSEGDGFAEDPTISLLTGSEPPKAKDPT
VS

**TasA-VPR:** SV40 Nuclear Localization Signal, *Flavonifractor plautii* TasA, VPR

PKKKRKVGIHGVPGGLEGGGGSGGTASMDIQKNRIRNIVGGIYDIQKLRIATGNRIVASLRP
GLVEEVKEGEEDTKYLPAILSEYRRITDYFVSEFEGRGSIEKAITPNNPEYIKSRLDYDLVTS
YKRLLETEEGLTKVAEREVKAHPMWDAFFAGVKGCGPLMSAVCLAYFDPYKARHASSFW
RYAGLDVQRDPDKDKMRGVGKWYTEERPYIDKDGKEQMKKSLTYNPFLKTKLVGVLGS
AFLRAKDSYYGKVYYDYKNRLDNRDEDFSPIVKHRMATRYAVKMFLRDMWVVWRELE
GLEVTEPYEVAKLGHKPHHSSSPKKKRKVGSEASGSGRADALDDFDLDMLGSDALDDFD
LDMLGSDALDDFDLDMLGSDALDDFDLDMLINSRSSGSPKKKRKVGSQYLPDTDDRHRI
EEKRKRTYETFKSIMKKSPFSGPTDPRPPPRRIAVPSRSSASVPKPAPQPYPFTSSLSTINYDE
FPTMVFPSGQISQASALAPAPPQVLPQAPAPAPAPAMVSALAQAPAPVPVLAPGPPQAVAPP
APKPTQAGEGTLSEALLQLQFDDEDLGALLGNSTDPAVFTDLASVDNSEFQQLLNQGIPVA
PHTTEPMLMEYPEAITRLVTGAQRPPDPAPAPLGAPGLPNGLLSGDEDFSSIADMDFSALL
GSGSGSRDSREGMFLPKPEAGSAISDVFEGREVCQPKRIRPFHPPGSPWANRPLPASLAPTP
TGPVHEPVGSLTPAPVPQPLDPAPAVTPEASHLLEDPDEETSQAVKALREMADTVIPQKEEA
AICGQMDLSHPPPRGHLDELTTTLESMTEDLNLDSPLTPELNEILDTFLNDECLLHAMHIST
GLSIFDTSLF

**TasA-10×GCN4:** SV40 Nuclear Localization Signal, *Flavonifractor plautii* TasA, 10×GCN4-GS linker

PKKKRKVGIHGVPGGLEGGGGSGGTASMDIQKNRIRNIVGGIYDIQKLRIATGNRIVASLRP
GLVEEVKEGEEDTKYLPAILSEYRRITDYFVSEFEGRGSIEKAITPNNPEYIKSRLDYDLVTS
YKRLLETEEGLTKVAEREVKAHPMWDAFFAGVKGCGPLMSAVCLAYFDPYKARHASSFW
RYAGLDVQRDPDKDKMRGVGKWYTEERPYIDKDGKEQMKKSLTYNPFLKTKLVGVLGS
AFLRAKDSYYGKVYYDYKNRLDNRDEDFSPIVKHRMATRYAVKMFLRDMWVVWRELE
GLEVTEPYEVAKLGHKPHHSSSPKKKRKVGSPKKKRKVNGPTDAAEEELLSKNYHLENE
VARLKKGSGSGEELLSKNYHLENEVARLKKGSGSGEELLSKNYHLENEVARLKKGSGSGE
ELLSKNYHLENEVARLKKGSGSGEELLSKNYHLENEVARLKKGSGSGEELLSKNYHLENE
VARLKKGSGSGEELLSKNYHLENEVARLKKGSGSGEELLSKNYHLENEVARLKKGSGSGE
ELLSKNYHLENEVARLKKGSGSGEELLSKNYHLENEVARLKKGSGSGTAVNIGGGTGPMD
LQPKKKRKV

**dTasR-XTEN80-NP:** SV40 Nuclear Localization Signal, *Parcubacteria* dTasR (D7A), XT80 linker, NP

PKKKRKVGIHGVPGGLEGGGGSGGTASMRQVAIAWAFAEEKLAVYDGKKVLKAVPKLKA
GDEIFAENIPMKHAAQWLKDGIIIRRCRPNDTAALRKEIGQEKTDALDAKLIWQLAEAHPE
KFREWKGDPQLTTLYRAFKEVQRCRVGQSNRVWAKGEETATAVLDDLEKTERKIVKAIEK

ELKSYKVWDWLSQIKGIGVATGGGLVGLIAKYGIENIRQVSSLWHLFGLHVVEGKAPRRT
KGEEVSYVPEAKTLVLGVIADCFIKQRSPVYRDIYDQEKARQLEIEYPEGELASKFLGYKK
SDTHLSLNHAHRRAIRKMMKIFVQHVWLAWRTCEGLETRPLYCHEYLGHEHFIEPPVKIQ
PEGSSPKKKRKVGGPSSGAPPPSGGSPAGSPTSTEEGTSESATPESGPGTSTEPSEGSAPGSP
AGSPTSTEEGTSTEPSEGSAPGTSTEPSEEGQSDERALLDQLHTLLSNTDATGLEEIDRALGI
PELVNQGQALEPKQGSGSGSHEKFPSDLDLDMFNGSLECDMESIIRSELMDADGLDFNFDS
GSGSGSFVTLKDVGMDFTLGDWEQLGLEQGDTFWDTALDNCQDLFLLGGGGSPSGQISN
QALALAPSSAPVLAQTMVPSSAMVPLAQPPAPAPVLTPGPPQSLSAPVPKSTQAGEGTLSE
ALLHLQFDADEDLGALLGNSTDPGVFTDLASVDNSEFQQLLNQGVSMSHSTAEPMLMEY
PEAITRLVTGSQRPPDPAPTPLGTSGLPNGLSGDEDFSSIADMDFSALLSQISSSGQGGGGSG
FSVDTSALLDLFSPSVTVPDMSLPDLDSSLASIQELLSPQEPPRPPEAENSSPDSGKQLVHYT
AQPLFLLDPGSVDTGSNDLPVLFELGEGSYFSEGDGFAEDPTISLLTGSEPPKAKDPTVS

**dTasR-XTEN80-VPR:** SV40 Nuclear Localization Signal, *Parcubacteria* dTasR (D7A), XT80 linker, VPR

PKKKRKVGIHGVPGGLEGGGGSGGTASMRQVAIAWAFAEEKLAVYDGKKVLKAVPKLKA
GDEIFAENIPMKHAAQWLKDGIIIRRCRPNDTAALRKEIGQEKTDALDAKLIWQLAEAHPE
KFREWKGDPQLTTLYRAFKEVQRCRVGQSNRVWAKGEETATAVLDDLEKTERKIVKAIEK
ELKSYKVWDWLSQIKGIGVATGGGLVGLIAKYGIENIRQVSSLWHLFGLHVVEGKAPRRT
KGEEVSYVPEAKTLVLGVIADCFIKQRSPVYRDIYDQEKARQLEIEYPEGELASKFLGYKK
SDTHLSLNHAHRRAIRKMMKIFVQHVWLAWRTCEGLETRPLYCHEYLGHEHFIEPPVKIQ
PEGSSPKKKRKVGGPSSGAPPPSGGSPAGSPTSTEEGTSESATPESGPGTSTEPSEGSAPGSP
AGSPTSTEEGTSTEPSEGSAPGTSTEPSEEASGSGRADALDDFDLDMLGSDALDDFDLDML
GSDALDDFDLDMLGSDALDDFDLDMLINSRSSGSPKKKRKVGSQYLPDTDDRHRIEEKR
KRTYETFKSIMKKSPFSGPTDPRPPPRRIAVPSRSSASVPKPAPQPYPFTSSLSTINYDEFPTM
VFPSGQISQASALAPAPPQVLPQAPAPAPAPAMVSALAQAPAPVPVLAPGPPQAVAPPAPKP
TQAGEGTLSEALLQLQFDDEDLGALLGNSTDPAVFTDLASVDNSEFQQLLNQGIPVAPHTT
EPMLMEYPEAITRLVTGAQRPPDPAPAPLGAPGLPNGLLSGDEDFSSIADMDFSALLGSGS
GSRDSREGMFLPKPEAGSAISDVFEGREVCQPKRIRPFHPPGSPWANRPLPASLAPTPTGPV
HEPVGSLTPAPVPQPLDPAPAVTPEASHLLEDPDEETSQAVKALREMADTVIPQKEEAAICG
QMDLSHPPPRGHLDELTTTLESMTEDLNLDSPLTPELNEILDTFLNDECLLHAMHISTGLSIF
DTSLF

**NP-dTasR:** SV40 Nuclear Localization Signal, NP, *Parcubacteria* dTasR (D7A)

PKKKRKVGIHGVPGGLEGGGGSEGQSDERALLDQLHTLLSNTDATGLEEIDRALGIPELVN
QGQALEPKQGSGSGSHEKFPSDLDLDMFNGSLECDMESIIRSELMDADGLDFNFDSGSGS
GSFVTLKDVGMDFTLGDWEQLGLEQGDTFWDTALDNCQDLFLLGGGGSPSGQISNQALA
LAPSSAPVLAQTMVPSSAMVPLAQPPAPAPVLTPGPPQSLSAPVPKSTQAGEGTLSEALLH
LQFDADEDLGALLGNSTDPGVFTDLASVDNSEFQQLLNQGVSMSHSTAEPMLMEYPEAIT
RLVTGSQRPPDPAPTPLGTSGLPNGLSGDEDFSSIADMDFSALLSQISSSGQGGGGSGFSVD
TSALLDLFSPSVTVPDMSLPDLDSSLASIQELLSPQEPPRPPEAENSSPDSGKQLVHYTAQPL
FLLDPGSVDTGSNDLPVLFELGEGSYFSEGDGFAEDPTISLLTGSEPPKAKDPTVSGGGGSG
GTASMRQVAIAWAFAEEKLAVYDGKKVLKAVPKLKAGDEIFAENIPMKHAAQWLKDGIII

RRCRPNDTAALRKEIGQEKTDALDAKLIWQLAEAHPEKFREWKGDPQLTTLYRAFKEVQ
RCRVGQSNRVWAKGEETATAVLDDLEKTERKIVKAIEKELKSYKVWDWLSQIKGIGVATG
GGLVGLIAKYGIENIRQVSSLWHLFGLHVVEGKAPRRTKGEEVSYVPEAKTLVLGVIADCF
IKQRSPVYRDIYDQEKARQLEIEYPEGELASKFLGYKKSDTHLSLNHAHRRAIRKMMKIFV
QHVWLAWRTCEGLETRPLYCHEYLGHEHFIEPPVKIQPEGSSPKKKRKV

**VPR-dTasR:** SV40 Nuclear Localization Signal, VPR, *Parcubacteria* dTasR (D7A)

PKKKRKVGIHGVPGGLEGGGGSEASGSGRADALDDFDLDMLGSDALDDFDLDMLGSDA
LDDFDLDMLGSDALDDFDLDMLINSRSSGSPKKKRKVGSQYLPDTDDRHRIEEKRKRTYE
TFKSIMKKSPFSGPTDPRPPPRRIAVPSRSSASVPKPAPQPYPFTSSLSTINYDEFPTMVFPSG
QISQASALAPAPPQVLPQAPAPAPAPAMVSALAQAPAPVPVLAPGPPQAVAPPAPKPTQAGE
GTLSEALLQLQFDDEDLGALLGNSTDPAVFTDLASVDNSEFQQLLNQGIPVAPHTTEPMLM
EYPEAITRLVTGAQRPPDPAPAPLGAPGLPNGLLSGDEDFSSIADMDFSALLGSGSGSRDSR
EGMFLPKPEAGSAISDVFEGREVCQPKRIRPFHPPGSPWANRPLPASLAPTPTGPVHEPVGS
LTPAPVPQPLDPAPAVTPEASHLLEDPDEETSQAVKALREMADTVIPQKEEAAICGQMDLS
HPPPRGHLDELTTTLESMTEDLNLDSPLTPELNEILDTFLNDECLLHAMHISTGLSIFDTSLF
GGGGSGGTASMRQVAIAWAFAEEKLAVYDGKKVLKAVPKLKAGDEIFAENIPMKHAAQW
LKDGIIIRRCRPNDTAALRKEIGQEKTDALDAKLIWQLAEAHPEKFREWKGDPQLTTLYRA
FKEVQRCRVGQSNRVWAKGEETATAVLDDLEKTERKIVKAIEKELKSYKVWDWLSQIKGI
GVATGGGLVGLIAKYGIENIRQVSSLWHLFGLHVVEGKAPRRTKGEEVSYVPEAKTLVLG
VIADCFIKQRSPVYRDIYDQEKARQLEIEYPEGELASKFLGYKKSDTHLSLNHAHRRAIRK
MMKIFVQHVWLAWRTCEGLETRPLYCHEYLGHEHFIEPPVKIQPEGSSPKKKRKV

**NP-XTEN80-dTasR:** SV40 Nuclear Localization Signal, NP, XT80 linker, *Parcubacteria* dTasR (D7A)

PKKKRKVGIHGVPGGLEGGGGSEGQSDERALLDQLHTLLSNTDATGLEEIDRALGIPELVN
QGQALEPKQGSGSGSHEKFPSDLDLDMFNGSLECDMESIIRSELMDADGLDFNFDSGSGS
GSFVTLKDVGMDFTLGDWEQLGLEQGDTFWDTALDNCQDLFLLGGGGSPSGQISNQALA
LAPSSAPVLAQTMVPSSAMVPLAQPPAPAPVLTPGPPQSLSAPVPKSTQAGEGTLSEALLH
LQFDADEDLGALLGNSTDPGVFTDLASVDNSEFQQLLNQGVSMSHSTAEPMLMEYPEAIT
RLVTGSQRPPDPAPTPLGTSGLPNGLSGDEDFSSIADMDFSALLSQISSSGQGGGGSGFSVD
TSALLDLFSPSVTVPDMSLPDLDSSLASIQELLSPQEPPRPPEAENSSPDSGKQLVHYTAQPL
FLLDPGSVDTGSNDLPVLFELGEGSYFSEGDGFAEDPTISLLTGSEPPKAKDPTVSGGPSSG
APPPSGGSPAGSPTSTEEGTSESATPESGPGTSTEPSEGSAPGSPAGSPTSTEEGTSTEPSEGS
APGTSTEPSEMRQVAIAWAFAEEKLAVYDGKKVLKAVPKLKAGDEIFAENIPMKHAAQWL
KDGIIIRRCRPNDTAALRKEIGQEKTDALDAKLIWQLAEAHPEKFREWKGDPQLTTLYRAF
KEVQRCRVGQSNRVWAKGEETATAVLDDLEKTERKIVKAIEKELKSYKVWDWLSQIKGIG
VATGGGLVGLIAKYGIENIRQVSSLWHLFGLHVVEGKAPRRTKGEEVSYVPEAKTLVLGVI
ADCFIKQRSPVYRDIYDQEKARQLEIEYPEGELASKFLGYKKSDTHLSLNHAHRRAIRKM
MKIFVQHVWLAWRTCEGLETRPLYCHEYLGHEHFIEPPVKIQPEGSSPKKKRKV

**VPR-XTEN80-dTasR:** SV40 Nuclear Localization Signal, VPR, XT80 linker, *Parcubacteria* dTasR (D7A)

PKKKRKVGIHGVPGGLEGGGGSEASGSGRADALDDFDLDMLGSDALDDFDLDMLGSDA
LDDFDLDMLGSDALDDFDLDMLINSRSSGSPKKKRKVGSQYLPDTDDRHRIEEKRKRTYE
TFKSIMKKSPFSGPTDPRPPPRRIAVPSRSSASVPKPAPQPYPFTSSLSTINYDEFPTMVFPSG
QISQASALAPAPPQVLPQAPAPAPAPAMVSALAQAPAPVPVLAPGPPQAVAPPAPKPTQAGE
GTLSEALLQLQFDDEDLGALLGNSTDPAVFTDLASVDNSEFQQLLNQGIPVAPHTTEPMLM
EYPEAITRLVTGAQRPPDPAPAPLGAPGLPNGLLSGDEDFSSIADMDFSALLGSGSGSRDSR
EGMFLPKPEAGSAISDVFEGREVCQPKRIRPFHPPGSPWANRPLPASLAPTPTGPVHEPVGS
LTPAPVPQPLDPAPAVTPEASHLLEDPDEETSQAVKALREMADTVIPQKEEAAICGQMDLS
HPPPRGHLDELTTTLESMTEDLNLDSPLTPELNEILDTFLNDECLLHAMHISTGLSIFDTSLF
GGPSSGAPPPSGGSPAGSPTSTEEGTSESATPESGPGTSTEPSEGSAPGSPAGSPTSTEEGTST
EPSEGSAPGTSTEPSEMRQVAIAWAFAEEKLAVYDGKKVLKAVPKLKAGDEIFAENIPMKH
AAQWLKDGIIIRRCRPNDTAALRKEIGQEKTDALDAKLIWQLAEAHPEKFREWKGDPQLT
TLYRAFKEVQRCRVGQSNRVWAKGEETATAVLDDLEKTERKIVKAIEKELKSYKVWDWL
SQIKGIGVATGGGLVGLIAKYGIENIRQVSSLWHLFGLHVVEGKAPRRTKGEEVSYVPEAK
TLVLGVIADCFIKQRSPVYRDIYDQEKARQLEIEYPEGELASKFLGYKKSDTHLSLNHAHR
RAIRKMMKIFVQHVWLAWRTCEGLETRPLYCHEYLGHEHFIEPPVKIQPEGSSPKKKRKV

**NP-tTasR:** SV40 Nuclear Localization Signal, NP, truncated *Parcubacteria* TasR

PKKKRKVGIHGVPGGLEGGGGSEGQSDERALLDQLHTLLSNTDATGLEEIDRALGIPELVN
QGQALEPKQGSGSGSHEKFPSDLDLDMFNGSLECDMESIIRSELMDADGLDFNFDSGSGS
GSFVTLKDVGMDFTLGDWEQLGLEQGDTFWDTALDNCQDLFLLGGGGSPSGQISNQALA
LAPSSAPVLAQTMVPSSAMVPLAQPPAPAPVLTPGPPQSLSAPVPKSTQAGEGTLSEALLH
LQFDADEDLGALLGNSTDPGVFTDLASVDNSEFQQLLNQGVSMSHSTAEPMLMEYPEAIT
RLVTGSQRPPDPAPTPLGTSGLPNGLSGDEDFSSIADMDFSALLSQISSSGQGGGGSGFSVD
TSALLDLFSPSVTVPDMSLPDLDSSLASIQELLSPQEPPRPPEAENSSPDSGKQLVHYTAQPL
FLLDPGSVDTGSNDLPVLFELGEGSYFSEGDGFAEDPTISLLTGSEPPKAKDPTVSGGGGSD
PQLTTLYRAFKEVQRCRVGQSNRVWAKGEETATAVLDDLEKTERKIVKAIEKELKSYKVW
DWLSQIKGIGVATGGGLVGLIAKYGIENIRQVSSLWHLFGLHVVEGKAPRRTKGEEVSYVP
EAKTLVLGVIADCFIKQRSPVYRDIYDQEKARQLEIEYPEGELASKFLGYKKSDTHLSLNH
AHRRAIRKMMKIFVQHVWLAWRTCEGLETRPLYCHEYLGHEHFIEPPVKIQPEGSSPKKK
RKV

**VPR-tTasR:** SV40 Nuclear Localization Signal, VPR, truncated *Parcubacteria* TasR

PKKKRKVGIHGVPGGLEGGGGSEASGSGRADALDDFDLDMLGSDALDDFDLDMLGSDA
LDDFDLDMLGSDALDDFDLDMLINSRSSGSPKKKRKVGSQYLPDTDDRHRIEEKRKRTYE
TFKSIMKKSPFSGPTDPRPPPRRIAVPSRSSASVPKPAPQPYPFTSSLSTINYDEFPTMVFPSG
QISQASALAPAPPQVLPQAPAPAPAPAMVSALAQAPAPVPVLAPGPPQAVAPPAPKPTQAGE
GTLSEALLQLQFDDEDLGALLGNSTDPAVFTDLASVDNSEFQQLLNQGIPVAPHTTEPMLM
EYPEAITRLVTGAQRPPDPAPAPLGAPGLPNGLLSGDEDFSSIADMDFSALLGSGSGSRDSR

EGMFLPKPEAGSAISDVFEGREVCQPKRIRPFHPPGSPWANRPLPASLAPTPTGPVHEPVGS
LTPAPVPQPLDPAPAVTPEASHLLEDPDEETSQAVKALREMADTVIPQKEEAAICGQMDLS
HPPPRGHLDELTTTLESMTEDLNLDSPLTPELNEILDTFLNDECLLHAMHISTGLSIFDTSLF
GGGGSDPQLTTLYRAFKEVQRCRVGQSNRVWAKGEETATAVLDDLEKTERKIVKAIEKEL
KSYKVWDWLSQIKGIGVATGGGLVGLIAKYGIENIRQVSSLWHLFGLHVVEGKAPRRTKG
EEVSYVPEAKTLVLGVIADCFIKQRSPVYRDIYDQEKARQLEIEYPEGELASKFLGYKKSD
THLSLNHAHRRAIRKMMKIFVQHVWLAWRTCEGLETRPLYCHEYLGHEHFIEPPVKIQPE
GSSPKKKRKV

**NP-XTEN80-tTasR:** SV40 Nuclear Localization Signal, NP, XT80 linker, truncated *Parcubacteria* TasR

PKKKRKVGIHGVPGGLEGGGGSEGQSDERALLDQLHTLLSNTDATGLEEIDRALGIPELVN
QGQALEPKQGSGSGSHEKFPSDLDLDMFNGSLECDMESIIRSELMDADGLDFNFDSGSGS
GSFVTLKDVGMDFTLGDWEQLGLEQGDTFWDTALDNCQDLFLLGGGGSPSGQISNQALA
LAPSSAPVLAQTMVPSSAMVPLAQPPAPAPVLTPGPPQSLSAPVPKSTQAGEGTLSEALLH
LQFDADEDLGALLGNSTDPGVFTDLASVDNSEFQQLLNQGVSMSHSTAEPMLMEYPEAIT
RLVTGSQRPPDPAPTPLGTSGLPNGLSGDEDFSSIADMDFSALLSQISSSGQGGGGSGFSVD
TSALLDLFSPSVTVPDMSLPDLDSSLASIQELLSPQEPPRPPEAENSSPDSGKQLVHYTAQPL
FLLDPGSVDTGSNDLPVLFELGEGSYFSEGDGFAEDPTISLLTGSEPPKAKDPTVSGGPSSG
APPPSGGSPAGSPTSTEEGTSESATPESGPGTSTEPSEGSAPGSPAGSPTSTEEGTSTEPSEGS
APGTSTEPSEDPQLTTLYRAFKEVQRCRVGQSNRVWAKGEETATAVLDDLEKTERKIVKAI
EKELKSYKVWDWLSQIKGIGVATGGGLVGLIAKYGIENIRQVSSLWHLFGLHVVEGKAPR
RTKGEEVSYVPEAKTLVLGVIADCFIKQRSPVYRDIYDQEKARQLEIEYPEGELASKFLGY
KKSDTHLSLNHAHRRAIRKMMKIFVQHVWLAWRTCEGLETRPLYCHEYLGHEHFIEPPVK
IQPEGSSPKKKRKV

**VPR-XTEN80-tTasR:** SV40 Nuclear Localization Signal, VPR, XT80 linker, truncated *Parcubacteria* TasR

PKKKRKVGIHGVPGGLEGGGGSEASGSGRADALDDFDLDMLGSDALDDFDLDMLGSDA
LDDFDLDMLGSDALDDFDLDMLINSRSSGSPKKKRKVGSQYLPDTDDRHRIEEKRKRTYE
TFKSIMKKSPFSGPTDPRPPPRRIAVPSRSSASVPKPAPQPYPFTSSLSTINYDEFPTMVFPSG
QISQASALAPAPPQVLPQAPAPAPAPAMVSALAQAPAPVPVLAPGPPQAVAPPAPKPTQAGE
GTLSEALLQLQFDDEDLGALLGNSTDPAVFTDLASVDNSEFQQLLNQGIPVAPHTTEPMLM
EYPEAITRLVTGAQRPPDPAPAPLGAPGLPNGLLSGDEDFSSIADMDFSALLGSGSGSRDSR
EGMFLPKPEAGSAISDVFEGREVCQPKRIRPFHPPGSPWANRPLPASLAPTPTGPVHEPVGS
LTPAPVPQPLDPAPAVTPEASHLLEDPDEETSQAVKALREMADTVIPQKEEAAICGQMDLS
HPPPRGHLDELTTTLESMTEDLNLDSPLTPELNEILDTFLNDECLLHAMHISTGLSIFDTSLF
GGPSSGAPPPSGGSPAGSPTSTEEGTSESATPESGPGTSTEPSEGSAPGSPAGSPTSTEEGTST
EPSEGSAPGTSTEPSEDPQLTTLYRAFKEVQRCRVGQSNRVWAKGEETATAVLDDLEKTER
KIVKAIEKELKSYKVWDWLSQIKGIGVATGGGLVGLIAKYGIENIRQVSSLWHLFGLHVVE
GKAPRRTKGEEVSYVPEAKTLVLGVIADCFIKQRSPVYRDIYDQEKARQLEIEYPEGELAS
KFLGYKKSDTHLSLNHAHRRAIRKMMKIFVQHVWLAWRTCEGLETRPLYCHEYLGHEHF
IEPPVKIQPEGSSPKKKRKV

**Table S2. Spacer sequences of sgRNAs used in Cas9-mediated homology directed repair for EGFP knock-in.**

| **Gene** | **Spacer sequences (5'-3')** |
|---|---|
| *RHOXF2* | TCTCATCCCGCTGATGAACA |
| *NEUROG2* | GGTGCATAGCGGTGCTTGTC |

**Table S3. The tigRNA guide sequences used in this study.**

| Gene | Spacer 1 sequence (5'-3') | Spacer 2 sequence (5'-3') |
|---|---|---|
| *RHOXF2*[1] | TasA tig1: ACGCGTGCTC | GATGAGGGA |
| | TasA tig2: CGCGTGCTCT | GGATGAGGG |
| | TasA tig3: CATCCTACTC | TGGGAGGGG |
| | TasA tig4: GGCCCAAGCT | AGGAGCAGG |
| | TasR tig1: CGCGTGCTC | GATGAGGGA |
| | TasR tig2: GCGTGCTCT | GGATGAGGG |
| | TasR tig3: CATCCTACT | GGGAGGGGG |
| | TasR tig4: GGCCCAAGC | GGAGCAGGA |
| *NEUROG2*[1] | TasA tig1: GCGGTGGCGG | CCTCCTCCC |
| | TasA tig2: AATGAAAAGA | CTGGCTTAT |
| | TasA tig3: GGAAAGGCGG | CTTTCTTCA |
| | TasA tig4: CTGCGGCTTC | GGAGCTGGC |
| | TasR tig1: CGGTGGCGG | CCTCCTCCC |
| | TasR tig2: ATGAAAAGA | CTGGCTTAT |
| | TasR tig3: GAAAGGCGG | CTTTCTTCA |
| | TasR tig4: CTGCGGCTT | GAGCTGGCG |
| *HBG*[2] | TasA tig1: GCTAGGGATG | AAGAATAAA |
| | TasA tig2: TTGACCAATA | GCCTTGACA |
| | TasA tig3: TGCAAATATC | TGTCTGAAA |
| | TasA tig4: AAATTAGCAG | TATCCTCTT |
| | TasR tig1: CTAGGGATG | AAGAATAAA |
| | TasR tig2: TGACCAATA | GCCTTGACA |
| | TasR tig3: GCAAATATC | TGTCTGAAA |
| | TasR tig4: AATTAGCAG | TATCCTCTT |
| *ASCL1*[1] | TasA tig1: GGGAGAAAGG | AACGGGAGG |
| | TasA tig2: AGAACTTGAA | GCAAAGCGC |
| | TasA tig3: CCAATTTCTA | GGGTCACCG |

| | | |
|---|---|---|
| | TasA tig4: TTGTGAGCCG | TCCTGTAGG |
| | TasR tig1: GGAGAAAGG | AACGGGAGG |
| | TasR tig2: GAACTTGAA | GCAAAGCGC |
| | TasR tig3: CAATTTCTA | GGGTCACCG |
| | TasR tig4: TGTGAGCCG | TCCTGTAGG |
| *TTN*[1] | TasR tig1: TTGGTGAAG | CAAAGGAGA |
| | TasR tig2: GTTAAAATC | GCATTTTCG |
| | TasR tig3: GCACAGTCC | CAAACCTGA |
| | TasR tig4: GAGCTCTCT | TAACGTTGA |
| *IL1B*[3] | TasR tig1: TAAACTGAG | GAGAATTAT |
| | TasR tig2: AACTGCACA | GACAATCGT |
| | TasR tig3: TTCTTTGTA | AAACTTAAG |
| | TasR tig4: CACACCCTC | GTCTGTATT |
| *IL1RN*[4,5] | TasR tig1: ATGCCAAGC | ACTGGGCCT |
| | TasR tig2: GTTTCTGCT | ACTCAGGGCT |
| | TasR tig3: TACTCTCTG | AGAGCACCT |
| | TasR tig4: TCTGCGTAA | CCTCCCATC |
| *CD2*[5] | TasR tig1: CATCTTTTA | ATGTTACTG |
| | TasR tig2: TGTTACTGT | ACATCTTTT |
| | TasR tig3: CCTATATTT | ACCACATAG |
| | TasR tig4: GCTTCTTGT | CTTTTGTAA |
| *IFNG*[5] | TasR tig1: AGATGAGAT | TCTGTCACC |
| | TasR tig2: TATTAATAA | AAACCTTAG |
| | TasR tig3: TACCTCCCC | GGGCGAAGT |
| | TasR tig4: CCAGGGCGA | CTCCCCACT |
| *HBB*[6,7] | TasR tig1: AAGAGCCAA | TACCTGTCC |
| | TasR tig2: GACAGCCGT | AGGACAGGT |
| | TasR tig3: CTCTTCTGG | AAGCCAGTG |
| | TasR tig4: TCTTAGAGG | TCAGCCCTC |

| | | |
|---|---|---|
| *MYOD1*[8] | TasR tig1: CTGGGCTCC | AAACGCCCC |
| | TasR tig2: GCCCCTGCG | CGGGGTGGC |
| | TasR tig3: TCCCTCCCT | CTACCGGGC |
| | TasR tig4:GGTTTGGAA | GCACGCCCT |
| *CYC1p-* | TasR tig1: ACTTTAGTG | ATGTGTCAG |
| *mCherry*[9] | TasR tig2: GCGTGTATA | CACGCTATA |
| | TasR tig3: GGATGGCCA | AAAGTTGCC |
| | TasR tig4: GCCAGGCGT | TATATATAC |
| | TasR tig5: GTGTATATA | TCCACGACT |
| | TasR tig6: AAGACCAAA | AACTGGCGC |

**Table S4. The qRT-PCR primers used in this study.**

| Gene | Primers (5'-3') | Reference |
|---|---|---|
| *HBG* | F: GCTGAGTGAACTGCACTGTGA<br>R: GAATTCTTTGCCGAAATGGA | 5 |
| *ASCL1* | F: GGGCTCTTACGACCCGCTCA<br>R: AGGTTGTGCGATCACCCTGCTT | 5 |
| *TTN* | F: TGTTGCCACTGGTGCTAAAG<br>R: ACAGCAGTCTTCTCCGCTTC | 1 |
| *IL1B* | F: ATGATGGCTTATTACAGTGGCAA<br>R: GTCGGAGATTCGTAGCTGGA | 3 |
| *IL1RN* | F: GGAATCCATGGAGGGAAGAT<br>R: TGTTCTCGCTCAGGTCAGTG | 10 |
| *CD2* | F: GTCAGCAAGGAATCCAGTGTCG<br>R: AACGAGCAGTGCCACAAAGACC | 10 |
| *IFNG* | F: GAGTGTGGAGACCATCAAGGA<br>R: TGTATTGCTTTGCGTTGGAC | 11 |
| *HBB* | F: GCACGTGGATCCTGAGAACT<br>R: ATTGGACAGCAAGAAAGCGAG | 5 |
| *MYOD1* | F: CTCCAACTGCTCCGACGGCAT<br>R: AGGCAGTCTAGGCTCGACAC | 11 |
| *GAPDH* | F: CAATGACCCCTTCATTGACC<br>R: TTGATTTTGGAGGGATCTCG | 5 |
